# FAST-MaP: Chemical Mapping of RNA Structures Using Primer-less Sequencing

**DOI:** 10.64898/2026.09.22.753544

**Authors:** Jigyasa Verma, Hamish M. Blair, Wipapat Kladwang, Rhiju Das

## Abstract

RNA structure governs the function of non-coding RNAs and influences mRNA stability and translation, making testing of models of experimental RNA structure relevant to fields spanning structural biology, virology, and molecular therapeutics. Here we present FAST-MaP (Fast and Accessible Sequencing Technology for Mutational Profiling), a protocol that enables per-nucleotide RNA structure characterization using standard molecular biology equipment and requiring no sequencing infrastructure or bioinformatics expertise. RNA is chemically modified with orthogonal probes (2A3 and DMS), reverse-transcribed, and PCR-amplified to produce dsDNA amplicons that are submitted directly to a commercial primer-less sequencing service. Returned FASTQ files are processed through a freely accessible web server to generate normalized reactivity profiles within minutes. The complete protocol, from DNA template to structural data, can be completed in approximately one week. We illustrate the workflow on a 659-nucleotide RNA, demonstrating how to test structure preservation across buffers, and how to test specific secondary and tertiary structure predictions of the RNA from computational modeling or cryo-electron microscopy. The protocol requires only standard molecular-biology skills and does not require sequencing or bioinformatics expertise.

## INTRODUCTION

RNA structure underlies RNA function, yet identifying which nucleotides are base-paired, flexible, or solvent-accessible remains experimentally difficult for most laboratories. Per-nucleotide structural information is useful across fields from structural biology to virology to RNA therapeutics, but the methods that provide it are not equally accessible to all groups. High-resolution methods such as cryo-electron microscopy (cryo-EM), nuclear magnetic resonance (NMR) spectroscopy, and X-ray crystallography can determine three-dimensional RNA structures, but they require specialized infrastructure, extensive technical expertise, and substantial quantities of purified, conformationally homogeneous RNA. Chemical probing offers a more accessible alternative. Methods such as DMS-MaPseq^1^, SHAPE-MaP^2^, and icSHAPE^3^ use small-molecule reagents to modify RNA at unpaired or flexible positions, then read out those modifications through sequencing, providing per-nucleotide structural information from microgram-scale samples. These methods have broadened access to RNA structural data but still require multiplexed NGS library preparation and bioinformatics expertise. For laboratories studying individual RNAs (to guide hypotheses, test predictions, screen cryo-EM candidates, or assess how variants, buffers, or ligands affect folding), the overhead of a complete chemical-probing pipeline remains prohibitive.

### Development of the protocol

Here we present FAST-MaP (Fast and Accessible Sequencing Technology for Mutational Profiling), a protocol that makes chemical probing accessible to laboratories without NGS library preparation or bioinformatics expertise. Current barriers to entry to chemical probing are not the chemical treatments, which are straightforward, but the surrounding steps of preparing and quality-controlling sequencing libraries, running the sequencers, and then demultiplexing and analyzing the data. Even simplified kits still require adapter ligation, indexing, and QC. FAST-MaP removes these barriers by outsourcing every step from library preparation through sequencing to commercial services.

FAST-MaP introduces no new chemistry; its modification, reverse transcription, and mutational-profiling steps draw on established methods^1,2,3,4^. Instead, it reorganizes the workflow around commercial primer-less services that accept individual dsDNA amplicons without user-supplied primers and return FASTQ files within days. Each probed RNA is converted to a dsDNA amplicon by reverse transcription and PCR and submitted directly as its own sample; reads span the full amplicon in one pass, so no adapter ligation, barcoding, or demultiplexing is required. The protocol uses two orthogonal probes: 2A3 acylates the 2′-OH of flexible nucleotides at all four bases, reporting backbone flexibility^4^, while DMS methylates unpaired adenines (N1-A) and cytosines (N3-C), reporting base-pairing status^1,5,6^. Because they interrogate different features, agreement confirms data quality while divergence reveals structure neither probe captures alone (see **Box 1**).

Analysis is performed with `cmuts` (count mutations; github.com/DasLab/cmuts), a C pipeline that aligns reads, counts per-nucleotide modifications, subtracts background against paired controls, and outputs normalized reactivity profiles^7^. For laboratories without computational support, the freely accessible `cmuts` web server automates the pipeline from uploaded FASTQ files, needing no local installation or command-line expertise. The full protocol runs from DNA template to reactivity profiles in about one week.

FAST-MaP was first applied by Townley et al.^7^ to chemically map c-di-GMP riboswitch designs, where it detected ligand-dependent structural changes at the binding site. Here we present the full protocol and extend it with a second orthogonal probe (2A3 alongside DMS), applying it to the 659-nt ROOL env-120 and using its per-nucleotide reactivity to test the RNA’s cryo-EM structural model. We also provide full experimental-design guidance, a step-by-step procedure, and troubleshooting, so the workflow is reproducible in labs without sequencing or bioinformatics infrastructure.

### Overview of the procedure

The workflow (**Fig. 1**) runs in seven stages from a synthetic-DNA template to a per-nucleotide reactivity profile: gene synthesis → prepare RNA → chemical modification → reverse transcription → cDNA PCR → sequencing service → reactivity profiles. These expand into 30 numbered steps in the Procedure. The procedure has three phases: sample preparation (Steps 1– 21; Days 1–2): transcription, folding, modification with 2A3 and DMS (each with a matched no-probe control), reverse transcription, and cDNA recovery; sequencing (Steps 22–26; Day 3): cDNA PCR, gel extraction, and submission of individual amplicons to a primer-less sequencing service; and analysis (Steps 27–30; Day 5): FASTQ files processed through `cmuts` to generate per-nucleotide reactivity profiles. End-to-end time is about one week, including a ∼6–48 h sequencing turnaround.

**Figure 1.**
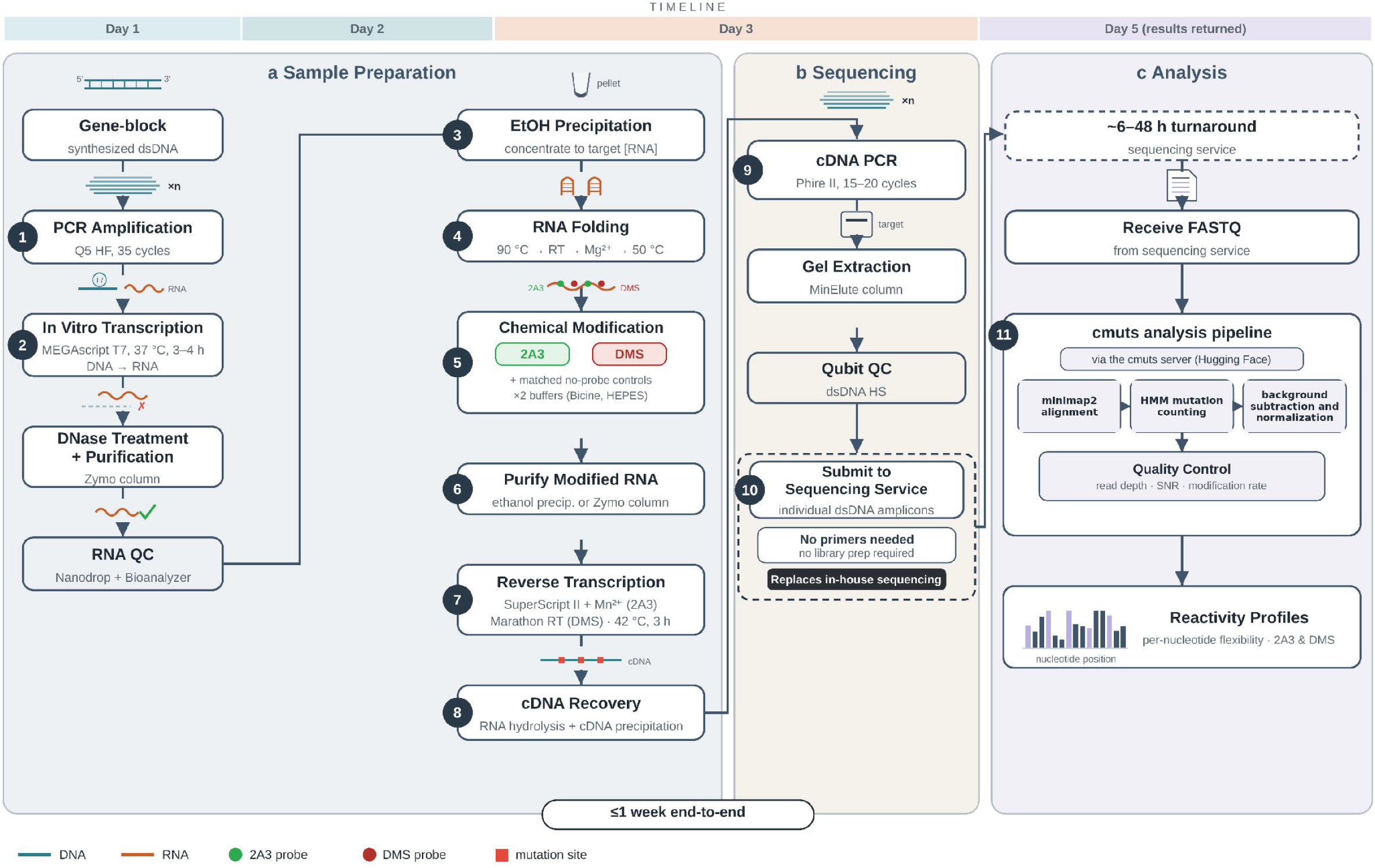
The FAST-MaP workflow. The full FAST-MaP workflow with intermediate steps, reagents, and color-coded biomolecules. (a) Sample preparation (Days 1–2): a DNA template is PCR-amplified from a synthetic gene fragment containing a T7 promoter, transcribed in vitro to produce RNA, assessed for concentration and integrity, and concentrated by ethanol precipitation. The RNA is folded under near-physiological conditions and chemically modified with either 2A3 or DMS, each with a matched no-probe control. Modified RNA is reverse-transcribed using probe-matched enzymes (SuperScript II with Mn2+ for 2A3; Marathon RT for DMS), and cDNA is recovered by alkaline hydrolysis and ethanol precipitation. (b) Sequencing submission (Day 3): cDNA is PCR-amplified, gel-extracted, quantified, and submitted as individual dsDNA amplicons to a commercial primer-less sequencing service. No primers or library preparation are required. (c) Analysis (Day 5): FASTQ files returned by the sequencing service (∼6–48 h turnaround) are processed through the FAST-MaP pipeline or the `cmuts` server to generate per-nucleotide reactivity profiles.

**Figure 2.**
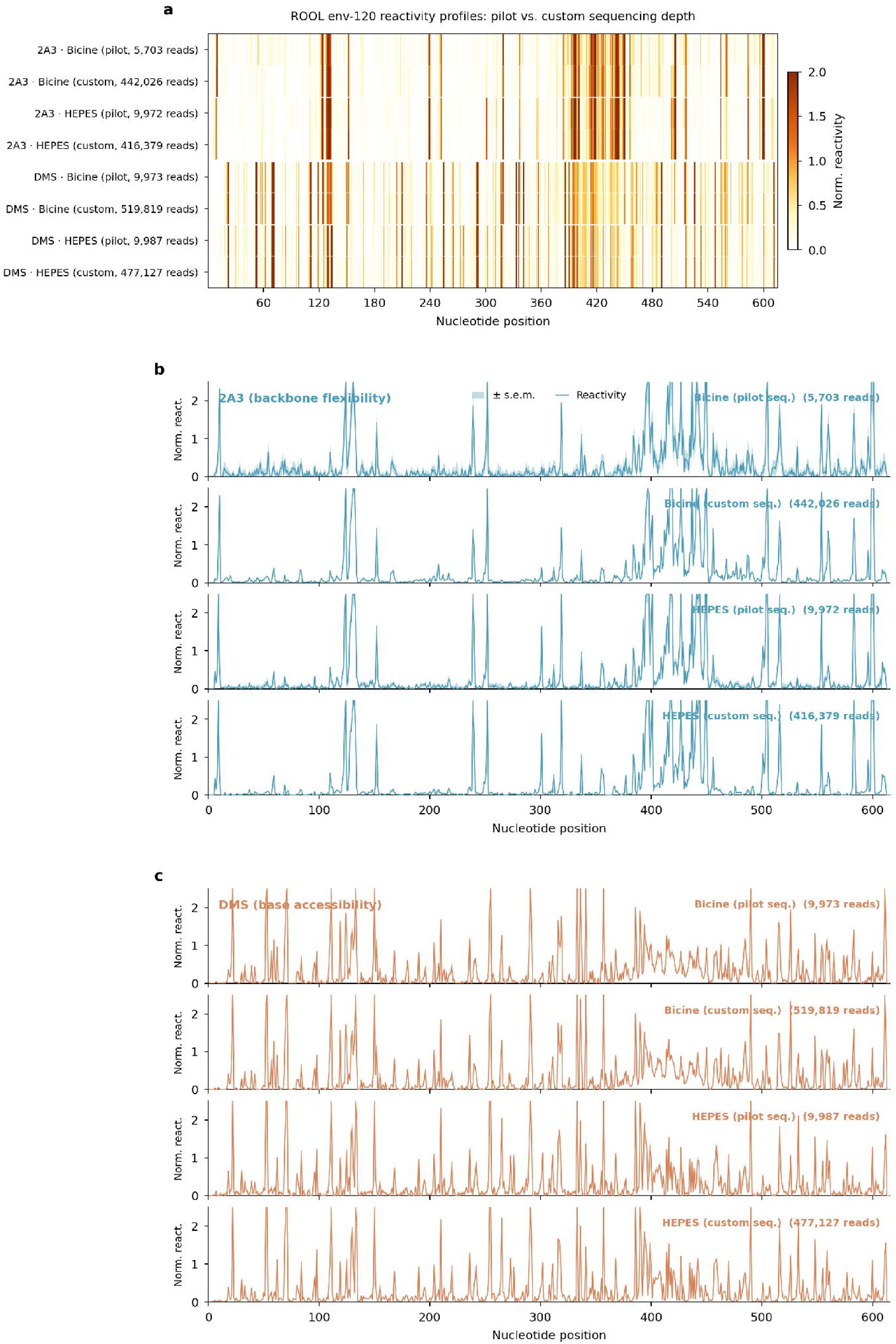
Per-nucleotide reactivity profiles for ROOL env-120 at pilot and custom sequencing depths. **(a)** Per-nucleotide reactivity heatmap across all 8 conditions (4 probe-buffer combinations × 2 sequencing depths). Rows are grouped by probe: 2A3 (Bicine pilot, Bicine custom, HEPES pilot, HEPES custom) then DMS (same order). Read counts are shown for each condition. **(b)** 2A3 reactivity line profiles with ±SE shaded bands, shown as four vertically stacked subplots matching the heatmap row order: Bicine pilot, Bicine custom, HEPES pilot, HEPES custom. The transition from pilot to custom sequencing shows substantially reduced SE bands. **(c)** DMS reactivity line profiles, same layout as (b). DMS achieves tighter SE bands than 2A3 at equivalent depths owing to the higher per-nucleotide signal of the DMS/Marathon RT system. Quantitative reproducibility scatter plots and sample-level SNR scaling are provided in **Supplementary Fig. 2**.

### Applications

FAST-MaP is designed for laboratories that need per-nucleotide structural information for one or a few RNAs at a time. A leading application is triage of computationally predicted and AI-designed structures^7^: reactivity ranks candidate models and flags mis-predicted regions before committing to cryo-EM. Other uses include screening for structural homogeneity before cryo-EM, assessing how variants affect folding, characterizing non-coding RNAs, and analyzing ligand- or protein-induced conformational changes. It also suits NGS-equipped laboratories wanting a rapid, low-overhead workflow. None of these use cases are tested here; most require only substituting the RNA of interest, although ligand- and protein-binding experiments may require condition-specific optimization. In the supporting study, for example, probing c-di-GMP riboswitch designs with and without ligand revealed localized reactivity protections at the binding site^7^. In-vivo probing with membrane-permeable reagents (e.g., in-cell DMS or SHAPE) is in principle compatible with the downstream workflow but is not tested here; for the probing step, consult DMS-MaPseq^1^, in-cell 2A3 probing^4^, and related protocols^3,5^.

### Comparison to other methods

FAST-MaP builds on existing RNA structure methods (**Table 1**). Compared with DMS-MaPseq^1^ and SHAPE-MaP^2^, it cuts workflow duration from 2–4 weeks to about one week. Its primary advantage is not per-sample pricing, which multiplexed NGS can drive to very low costs, but the elimination of workflow complexity: indexed library design, run management, and demultiplexing. For laboratories without NGS infrastructure this barrier is substantial; FAST-MaP replaces it with direct amplicon submission at tens of dollars per sample, with reagent costs comparable to other chemical-probing workflows.

**Table 1.**
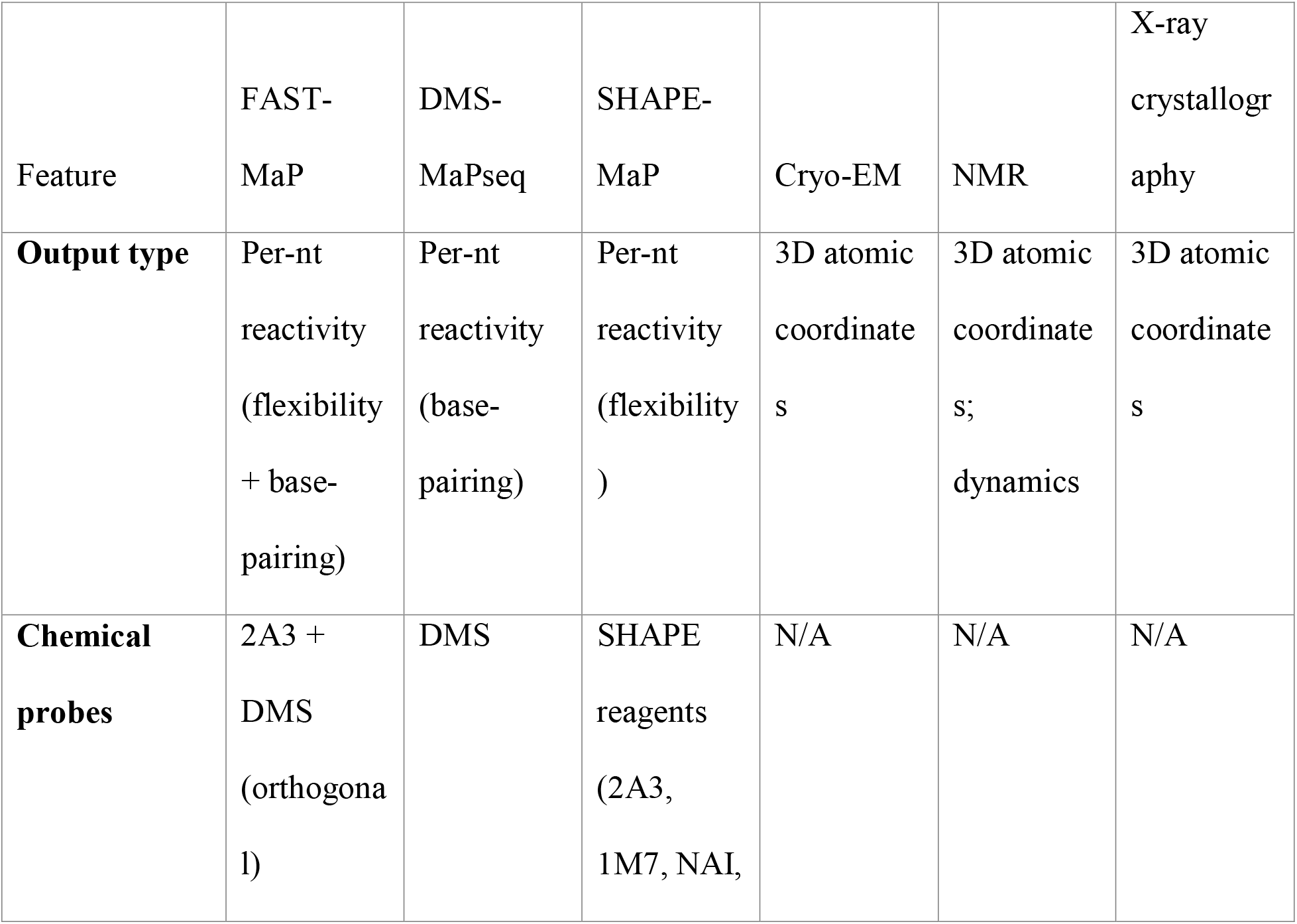
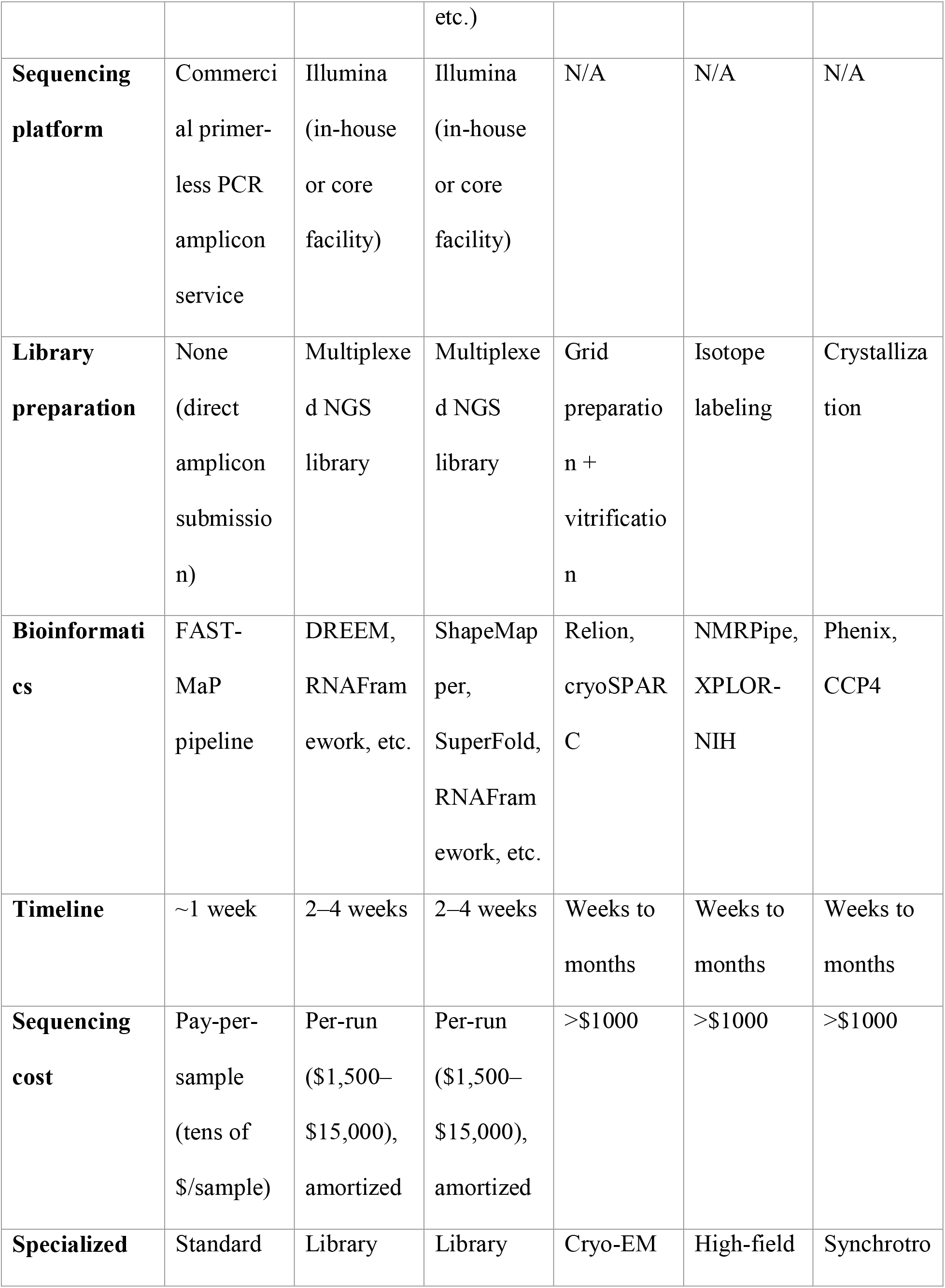
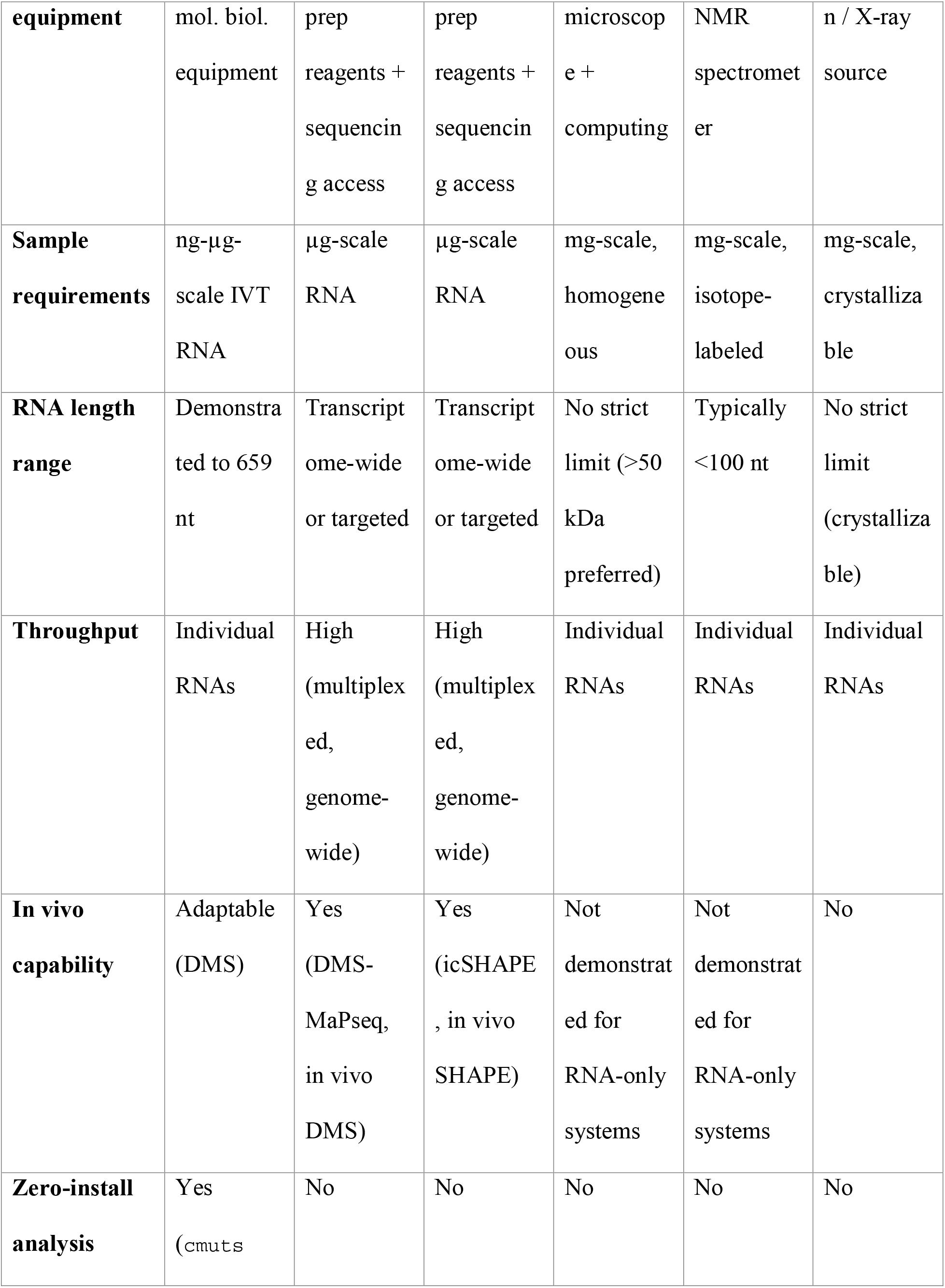
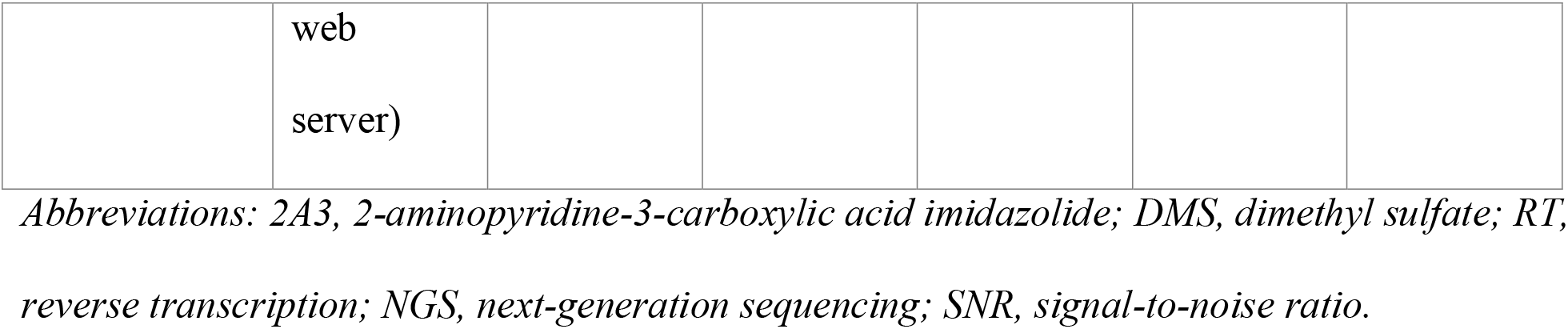
Comparison of FAST-MaP with established RNA structure determination methods.

Relative to high-resolution methods (cryo-EM, NMR, X-ray crystallography), FAST-MaP gives per-nucleotide flexibility and base-pairing information rather than atomic coordinates, a trade-off that makes it practical as a first-pass tool or a way to test computational predictions at a fraction of the cost and time.

FAST-MaP is vendor-agnostic, requires no kit purchase, and offers a zero-installation analysis path through the `cmuts` web server. To our knowledge, no prior chemical-probing protocol has been designed and benchmarked specifically for commercial amplicon sequencing: user preparation ends at amplicon submission, and the sequencing provider handles library preparation and demultiplexing internally.

#### Limitations and future directions

Several limitations should be considered. First, FAST-MaP has been demonstrated on RNA lengths up to 659-nucleotides. We recommend FAST-MaP for RNAs of ∼100–700 nt and advise assessing RT efficiency empirically for longer targets.

Second, pilot sequencing gave usable DMS profiles (SNR ≈ 3.6–4.5) but weaker 2A3 signal (SNR ≈ 1.2–2.5) for a 659-nt RNA. We therefore recommend DMS as the primary probe at low read depths, with 2A3 adding complementary backbone-flexibility information at custom depths (see Box 1); its moderate ROC AUC (0.723) reflects that it reports flexibility rather than base-pairing and is most informative alongside DMS (ROC AUC 0.815). Detecting subtle structural differences, e.g., upon addition of ligands or changes of buffer conditions, may require custom increased read depths.

Third, FAST-MaP produces population-averaged measurements; conformational heterogeneity is not directly resolved, though discordant 2A3/DMS reactivity can give indirect evidence for alternative conformations. Single-molecule deconvolution (e.g. DREEM^8^) could in principle be applied but requires high sequencing depths that are not by default available in commercial primer-less sequencing providers, and is untested. Fourth, the protocol uses two RT enzymes (SSII for 2A3, Marathon RT for DMS); single-probe users can simplify accordingly. Several directions could extend FAST-MaP. Improved direct-RNA Nanopore basecalling may eventually remove the RT and PCR steps, extending the length range and enabling single-molecule analysis; cheaper RNA synthesis could remove in vitro transcription; and membrane-permeable reagents such as DMS enable in vivo probing. Where services offer barcoded multiplexing, the method scales to higher throughput, and its reactivity can serve as restraints for computational structure prediction. As commercial sequencing costs fall, per-sample economics improve without changes to the wet-lab workflow.

### Experimental Design

#### RNA considerations

##### Target RNA suitability

FAST-MaP applies to any RNA that can be reverse-transcribed end-to-end. It is best suited where recovering secondary structure is the goal (riboswitches, ribozymes, regulatory elements, and functional domains of long non-coding RNAs), but also informs mRNAs, viral RNAs, and other less-structured targets. Highly repetitive or low-complexity sequences should be assessed empirically, as PCR slippage and truncated cDNA can degrade the reactivity profile.

The protocol is demonstrated here on ROOL env-120, a 659-nt structured RNA identified by comparative genomics of cow-rumen metagenomes^9^, whose three-dimensional structure has been independently determined by cryo-EM as an octameric nanocage at 3.1 Å resolution (PDB: 9MDS)^10^, a structural model that is tested by the FAST-MaP data.

##### Length range

FAST-MaP is bounded at the low end by the minimum insert commercial primer-less sequencing services accept (typically ∼100 nt) and at the high end by RT processivity. It is demonstrated here at 659 nt and would need to be assessed for longer RNAs empirically, since read-through depends on sequence and structure. For longer RNAs, reduce probe concentrations to limit over-modification that impedes full-length cDNA synthesis (see **Chemical probe considerations**, below). For transcripts longer than a single amplicon can cover by full-length reverse transcription and read-through, probe the full-length RNA once (using correspondingly higher RNA input), then amplify it as a series of overlapping windows with multiple primer pairs, with each window sized for reliable cDNA synthesis and within the sequencing service’s read length, and stitch the per-nucleotide reactivities across the shared overlaps during analysis. This tiling strategy is not demonstrated here but is well established for long and genome-scale RNAs (e.g. Smola et al.^11^; Manfredonia et al.^12^).

##### Template design

FAST-MaP probes RNA structure independent of how the RNA is produced. We demonstrate it with RNA from in vitro transcription (IVT) of a synthetic DNA template, which the procedure below follows. RNA from other sources (cell-extracted, enzymatic, or commercially prepared) can also be used and would enter FAST-MaP at the chemical-modification stage (Step 8) if it meets the purity and integrity criteria below, though we have not tested these. For IVT, a commercially synthesized gene fragment (e.g., IDT gBlocks or Twist Gene Fragments) encoding the RNA downstream of a T7 promoter (5’-TAATACGACTCACTATA-3’) is recommended, avoiding cloning. Efficient T7 initiation needs the transcript to begin with one or more guanosines (5’-GG or 5’-GGG); if it does not, add a short 5’-GG leader after the promoter. Most DNA synthesis providers impose length limits and reject high-repeat, homopolymer, or extreme-GC sequences; consult supplier guidelines and split long or GC-skewed targets into fragments joined by overlap PCR before IVT or consider specialized enzymatic synthesis of difficult templates.

Optionally, the gene fragment can include short 5′ and 3′ buffer regions flanking the RNA of interest (between the T7 promoter and the RNA at the 5′ end, and immediately downstream at the 3′ end) to serve as dedicated primer-binding sites (not used here). A primer pair annealing within these regions amplifies both the dsDNA template (Step 1) and the cDNA (Step 9), recovering every position of the RNA of interest. In the demonstration we probed the native ROOL env-120 sequence and did not add synthetic 5′ or 3′ buffer (flanking) regions. The forward and reverse primers therefore anneal at the RNA’s own 5′ and 3′ termini, over the first 20 and last 24 nucleotides, respectively. Because modifications within a primer footprint are overwritten during reverse transcription and PCR, these terminal nucleotides carry no reactivity, and reactivity is reported for the internal region of the transcript (positions 21 to 635, numbered 1 to 615 in the output). Studies needing reactivities at the extreme 5′ and 3′ ends should include dedicated 5′/3′ buffer regions so the primers anneal outside the RNA of interest or should capture the ends by adapter ligation to the RNA or to the cDNA^13,14,15^. Effective buffers are short, well-folded cassettes that fold independently of the target and do not base-pair with it, for example Das-lab GAGUA reference hairpins (e.g. 5′ GGAGACCUCGAGUAGAGGUCAAAA and 3′ AAACAACUCGAGUAGAGUUGACAAC), which also serve as internal reactivity standards^16^. To confirm a buffer is acceptable, fold the complete construct (buffers plus target) in a secondary-structure prediction program and check that the predicted base-pairing of the RNA of interest is unchanged relative to the target alone; if the buffer perturbs folding, redesign or lengthen it (or substitute a different reference hairpin) until the target folds independently.

##### Purity and quality control

RNA purity is critical. Remove residual DNA by DNase I digestion, verify A260/A280 ≥ 2.0 and A260/A230 ≥ 1.8 (preferably ≥ 2.0) after each purification, and confirm a single band or peak at the expected length by gel or capillary electrophoresis before chemical modification. Degraded RNA yields truncated cDNA and can lower final PCR amplicon yield or give rise to aberrant PCR products that contaminate the FAST-MaP signal.

#### Chemical probe considerations

##### Probe chemistry and readout

FAST-MaP uses two complementary probes: 2A3 reports on local 2′-OH dynamics at all four bases, DMS on Watson-Crick–face accessibility at adenosine and cytidine. See **Box 1** for guidance on selecting one or both probes based on experimental goals and resource constraints.

**2-aminopyridine-3-carboxylic acid imidazolide (2A3)**^4,17^ acylates the 2′-OH of conformationally flexible ribonucleotides (typically at unpaired positions) at all four ribonucleotides. The resulting 2′-O-acyl adduct is read out predominantly as a deletion by SuperScript II reverse transcriptase (SSII; used in the presence of Mn^2+^ throughout this protocol) with misincorporations occurring at a lower frequency^2,18^. The Mn^2+^ cofactor promotes read-through rather than stalling, essential for the deletion/mutation-profiling (MaP) readout.

**Dimethyl sulfate (DMS)**^1,5,6^ methylates the Watson-Crick face of adenosine (N1-A) and cytidine (N3-C) at unpaired, solvent-accessible positions. These methylations are read out predominantly as mutations by Marathon reverse transcriptase, a group II intron-derived enzyme optimized to read through methylated bases and incorporate a mismatch rather than stall^18,19^.

Other reverse transcriptases reported for DMS mutational profiling, including TGIRT and Ultramarathon, are expected to be compatible but are not tested here. DMS also methylates N7 of guanosine (at its Hoogsteen edge) at high frequency; because this methyl lies on the Hoogsteen face rather than the Watson–Crick face, N7-methylguanosine does not strongly block reverse transcription or depurinate under these conditions; instead, the N7-methylated base is prone to tautomerization that promotes nucleotide misincorporation (read predominantly as U) during reverse transcription^20^, so N7-G methylation contributes to the DMS profile; consistent with this, guanosines in the ROOL env-120 data show reactivity well above the unmodified-nucleotide (U) baseline and comparable to adenosine, indicating DMS reports at G as well as A and C under these conditions, although the signal reflects the environment of the G’s Hoogsteen edge and not on its Watson-Crick pairing status.

Because the probes report different chemistry, their profiles are not expected to correlate globally; agreement where both report protection or reactivity strengthens structural assignments. See the **CAUTION** flags in Materials and Procedure for DMS and 2A3 handling requirements.

##### Probe concentration and single-hit kinetics

For longer RNAs, probe working concentrations may need to be reduced to maintain single-hit modification kinetics (on average approximately one modification per RNA molecule)^21^.

In the experiments reported here, we used 133.3 mM 2A3 for the 659-nt ROOL env-120, approximately one-third of the 400 mM concentration recommended for shorter RNAs^4^. The same consideration applies to DMS: a 12% (v/v) DMS working stock giving 3% (v/v) final in the 20 µL reaction is typical for shorter RNAs^1,5^; for ROOL env-120 we reduced this to 4% (v/v) working / 1% (v/v) final (also about one-third) to limit multiple-hit events. Titrate both probes empirically for other lengths. RNA input is flexible and the high concentration here is not required: the demonstration used 4 µM final (80 pmol, 34 µM stock), but ≤1 µM straight from a spin-column (e.g. Zymo) eluate suffices for routine single-RNA probing, as is standard for SHAPE/DMS.

##### No-probe controls

Every modification reaction must be paired with a matched no-probe control (DMSO-only control for 2A3; ethanol-only control for DMS) processed identically through all downstream steps from modification through sequencing. These paired controls are used by `cmuts` for background subtraction of sequence-dependent RT errors, intrinsic deletion and mutations in the RNA, and errors from sequencing.

#### PCR and sequencing considerations

##### Amplicon purification and submission

Gel-extract the final amplicon to remove non-specific products and primer dimers. Verify concentration by fluorometry (e.g., Qubit dsDNA HS) or UV and confirm it meets the provider’s input requirements before submission (ROOL env-120: 7 ng/µL in 10 µL). Samples are submitted individually; the minimum experiment is one modified sample plus one matched no-probe control (2 samples), each incurring a separate charge.

##### Provider requirements

A compatible service must: (i) read the full amplicon in a single pass without requirement of primers (Oxford Nanopore); (ii) accept gel-extracted amplicons and perform all library preparation internally; (iii) return FASTQ files; and (iv) offer ∼48 h turnaround, with some pickup-based services achieving <6 h turnaround. All data here used Plasmidsaurus (plasmidsaurus.com), but the protocol is not dependent on this provider.

##### Read depth

We recommend a two-stage strategy: submit at the provider’s lowest-cost tier (typically 3000 reads per sample) to gauge quality (SNR of ∼2–3 or higher is usable), and if higher quality is desired, then place a custom order for higher sequencing read depth. For ROOL env-120, pilot submissions (∼5,700–10,000 reads) gave usable DMS profiles (SNR 3.6–4.5) but weaker 2A3 (SNR ≈ 1.2–2.5); a custom order (0.264 Gb, 8 samples) delivered ∼416,000– 520,000 reads per sample at SNR 14.0–29.8 for both probes (**Supplementary Fig. 2c**). For shorter RNAs, higher probe concentrations can achieve sufficient SNR at pilot sequencing depths without custom orders.

#### Controls and replication

##### No-probe controls

Every modification reaction must include a matched no-probe control (see Chemical probe considerations).

##### Replication

For uncertainty estimation, at least two independent biological replicates (separate IVT preps through the full protocol) are recommended. For ROOL env-120, cross-buffer reproducibility gave Pearson *r* = 0.895 (2A3) and *r* = 0.933 (DMS) (**Supplementary Fig. 2a**), and cross-depth reproducibility yielded *r* = 0.996 (DMS) and *r* = 0.980–0.998 (2A3) (**Supplementary Fig. 2b**).

#### Analysis setup

##### Pipeline overview

Reactivity profiles are generated using the `cmuts` (count mutations)^7^ pipeline. This aligns the FASTQ sequencing data to the reference library using `minimap2`, determines the location of probe-mediated errors, performs background subtraction, and optionally normalizes the final profiles. Processing takes under a minute per amplicon; the full walkthrough is in Step 11.

#### Reference FASTA preparation

The reference FASTA should be the full amplicon sequence, including the primer-binding regions at both ends. On the `cmuts` server, specify the forward- and reverse-primer lengths; `cmuts` reports reactivity for the internal region between the primers and assigns no reactivity (NaN) for the remainder, since modifications in the primer regions are overwritten during reverse transcription and PCR. Note the distinction between the reference you supply (the full amplicon, which sets the coordinate frame) and the positions that carry reactivity (the internal region): keep both in mind when using the profiles downstream, for example as input for machine-learning training, so reactivities map to the correct reference coordinates. In the ROOL env-120 demonstration the analysis reference was the internal region itself, transcript positions 21 to 635 (615 nucleotides, numbered 1 to 615 in the output), so every reported position carries reactivity and the primer-binding termini (transcript 1 to 20 and 636 to 659) were excluded..

##### Web server

The `cmuts` web server provides a browser-based interface that automates the full pipeline (**Fig. 3**). Users upload a reference FASTA and FASTQ files, choose normalization settings, and receive downloadable HDF5 and CSV outputs and interactive visualizations, with no local installation.

**Figure 3.**
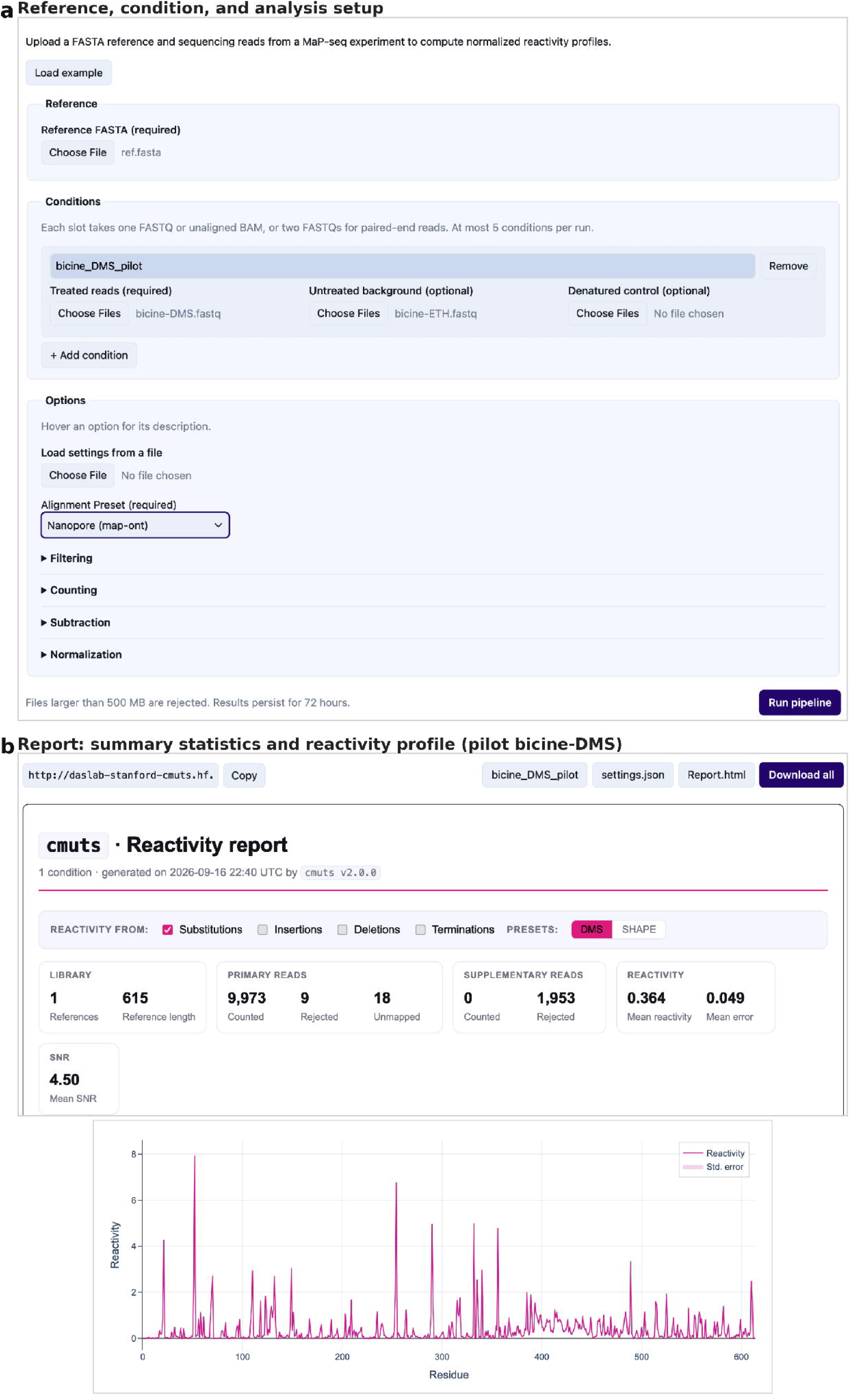
The cmuts server for zero-installation data analysis. **(a)** Reference, condition, and analysis setup. Users upload a reference FASTA and one or more experiment groups, each comprising a modified FASTQ and an optional control FASTQ; additional groups can be added to compare conditions, with normalization applied uniformly across all groups so values are directly comparable. Collapsible sections below the inputs hold the options for alignment, read filtering, counting, background subtraction, and normalization; sensible defaults require no tuning. **(b)** Example server output for ROOL env-120 (pilot DMS, bicine). The read depths, mean reactivity, mean error, and signal-to-noise ratio provide an at-a-glance summary of the quality of the data. The reactivity profile represents the sum of a configurable set of channels; two presets are tuned for SHAPE and DMS chemistry. Not pictured are per-base coverage plots and read length distributions for further quality control. The entire analysis can be performed within the browser without the need for local installation, with results persisting for 72 hours via a provided link.

#### Regulatory approvals

FAST-MaP requires no special regulatory approvals. It does not involve human or animal subjects, controlled substances, or select agents. All reagents, enzymes, and gene fragments are commercially available and handled under standard laboratory safety practice; users should confirm any material-transfer or licensing terms for their sources. The principal safety consideration is the chemical probe: dimethyl sulfate (DMS) is toxic and a suspected carcinogen and must be handled in a fume hood with appropriate personal protective equipment, with waste quenched and disposed of as hazardous chemical waste in accordance with institutional guidelines (see CAUTION callouts in Materials and Procedure).

## MATERIALS

### Reagents

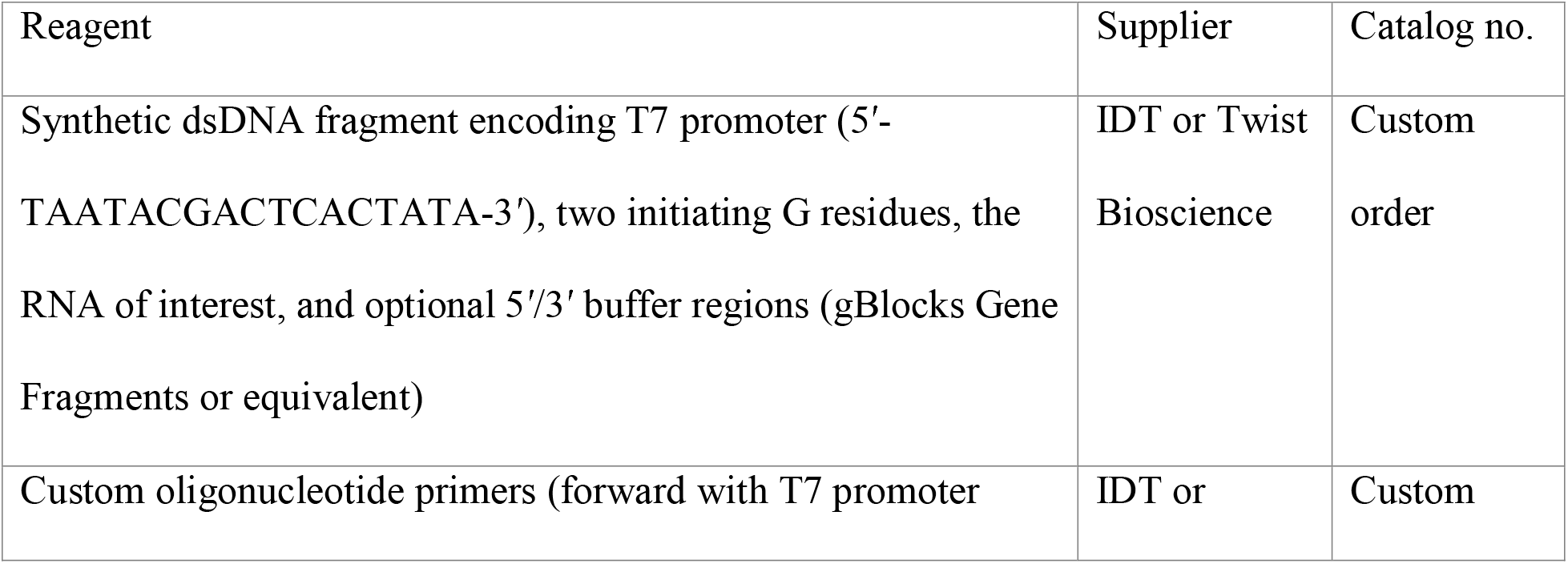
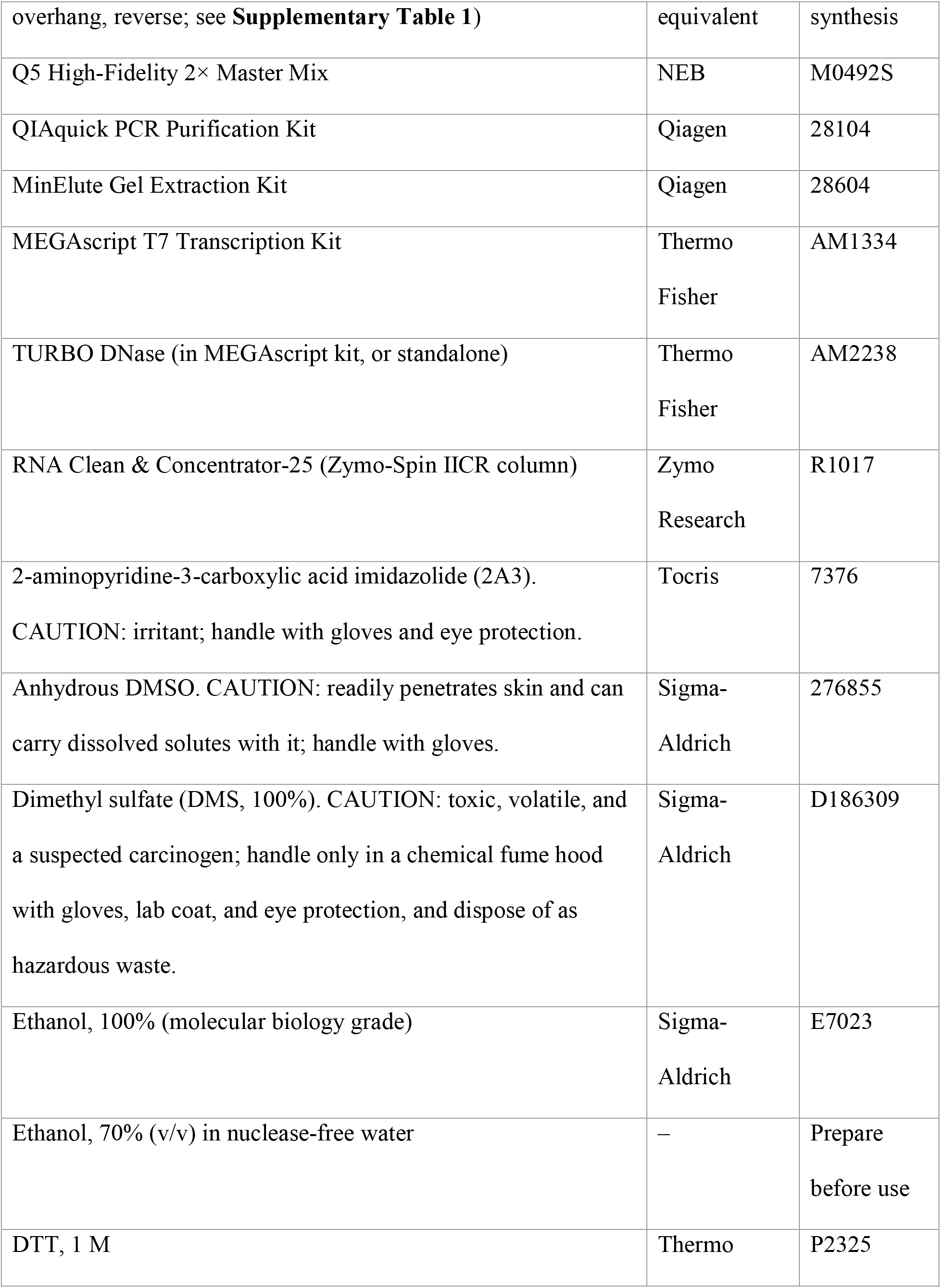
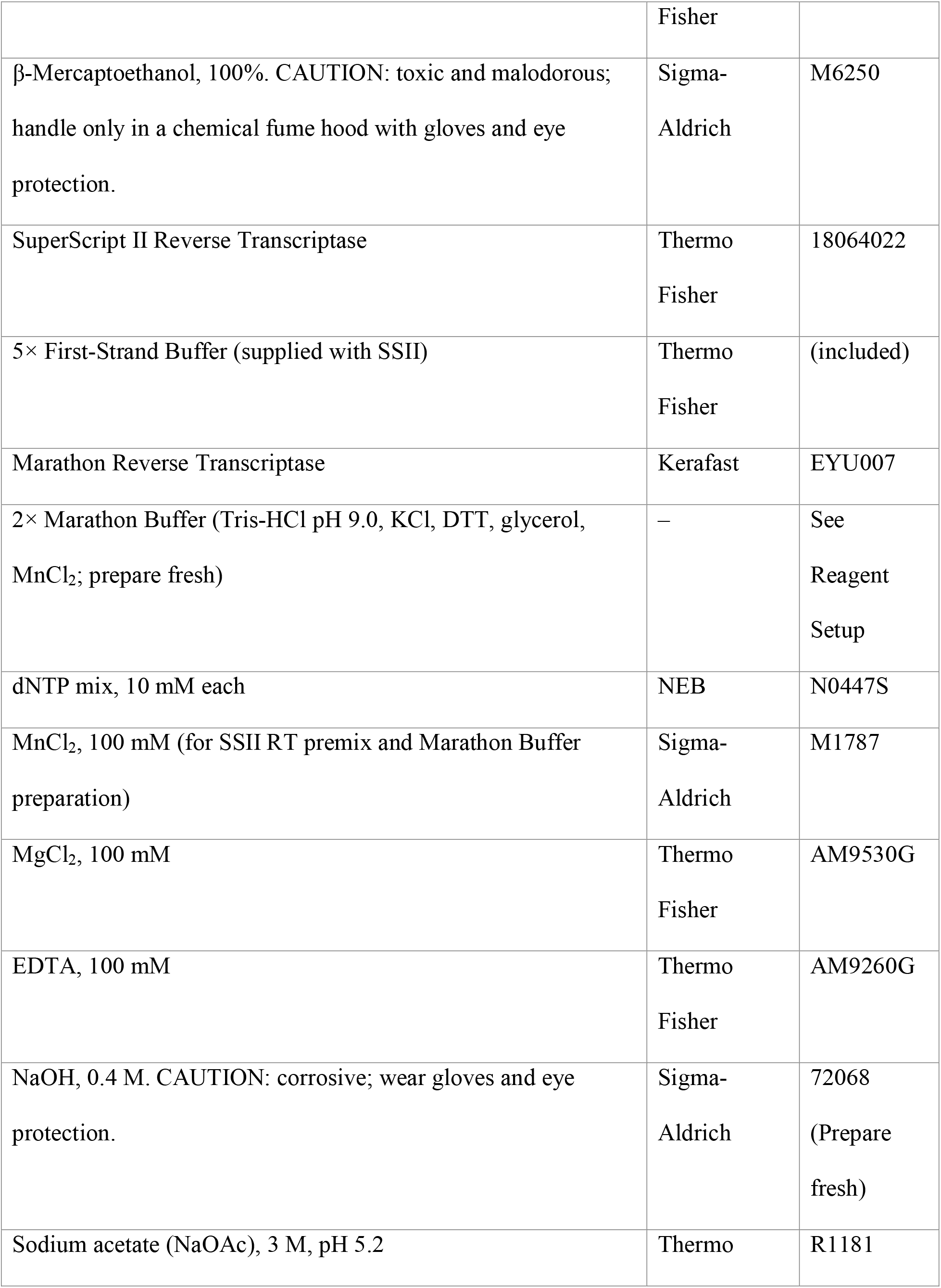
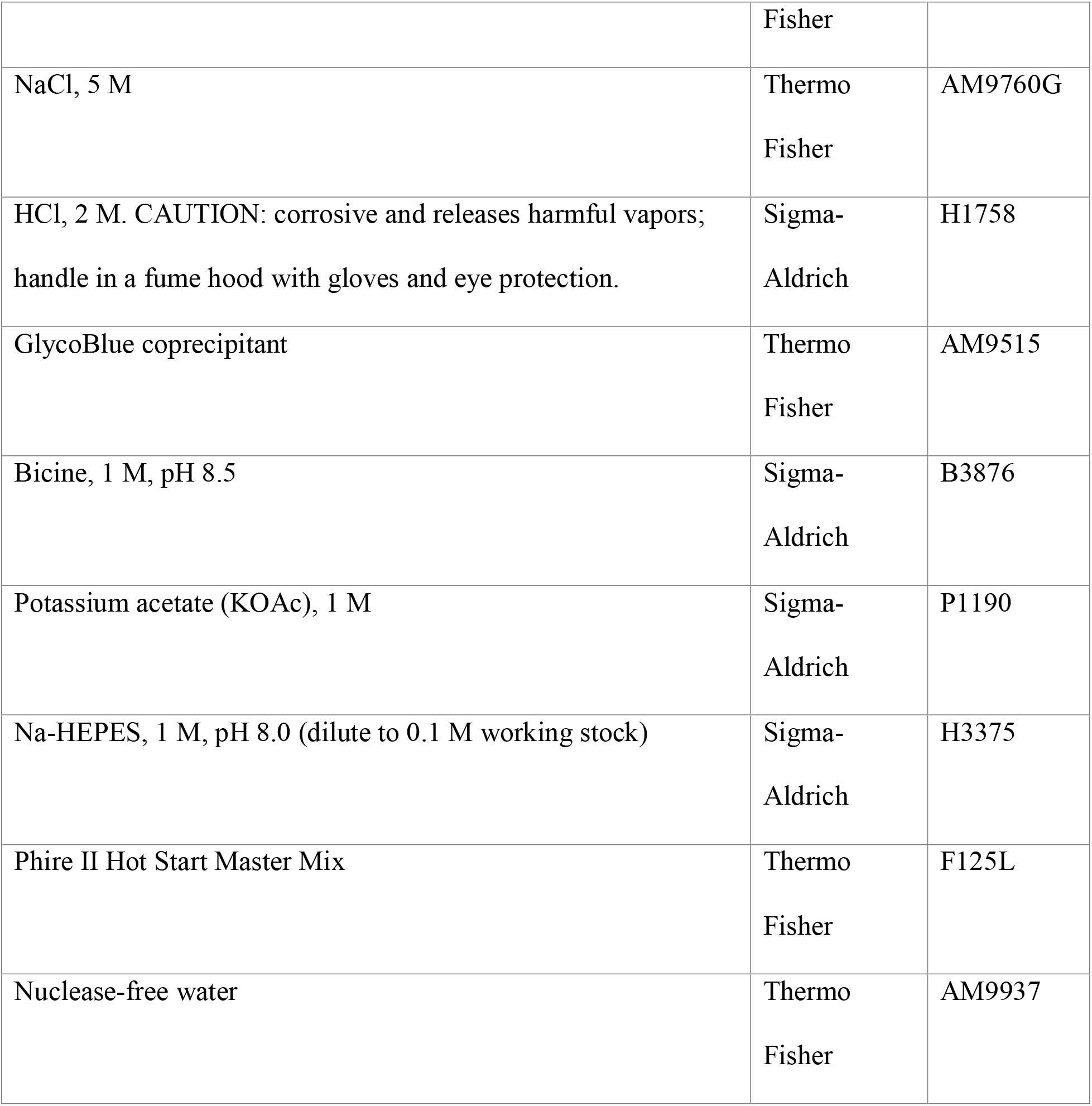

#### Optional reagents for enhanced QC (not required for the core protocol)

**Optional reagents for enhanced QC (not required for the core protocol):**
| Reagent | Supplier | Catalog no. |
| --- | --- | --- |
| Qubit dsDNA HS Assay Kit | Thermo Fisher | Q32851 |
| Qubit RNA HS Assay Kit | Thermo Fisher | Q32852 |
| Qubit ssDNA HS Assay Kit | Thermo Fisher | Q10212 |
| E-Gel EX or SYBR Safe 2% agarose cartridges | Thermo Fisher | G401002 |

**Note:** This protocol was demonstrated using two folding buffer conditions (Na-Bicine/KOAc at pH 8.5 and Na-HEPES at pH 8.0) as an optional test of whether the RNA structure changes across conditions; both yielded well-correlated reactivity profiles (see Controls and replication), so a single buffer condition is sufficient for routine applications. We do not recommend probing in two buffers routinely; because some RNAs can adopt buffer-dependent structures, however, probing in parallel conditions is informative when buffer-dependence is a concern or of scientific interest. Users may choose either buffer based on compatibility with their RNA system.

### Equipment

| Equipment | Supplier | Notes |
| --- | --- | --- |
| Thermocycler | Any | Standard PCR thermocycler; also used for RT and IVT incubations |
| Heat block (optional) | Any | Set to 37°C; alternative to thermocycler for IVT incubation |
| UV spectrophotometer or plate reader | Any | For DNA/RNA concentration by A260; any UV-capable instrument |
| Agarose gel electrophoresis system | Any | 1–2% gels with SYBR Safe or equivalent; E-Gel (Thermo) optional |
| Gel imaging system | Any | UV or blue-light transilluminator |
| Microcentrifuge (refrigerated) | Any | Capable of $\geq 16,000 \times g$ at 4°C |
| Chemical fume hood | – | Required for all DMS handling; not needed if using 2A3 only |
| Vortex mixer | Any |  |
| Pipettes and filtered tips | Any | 0.5–1,000 μL range |
| 1.5 mL microcentrifuge tubes | Any | Should be nuclease-free. DNA LoBind/low-retention tubes, e.g. Eppendorf 022431021, recommended for cDNA storage. |
| PCR tubes or strip tubes | Any | 0.2 mL, nuclease-free |

#### Optional equipment for enhanced QC (not required)

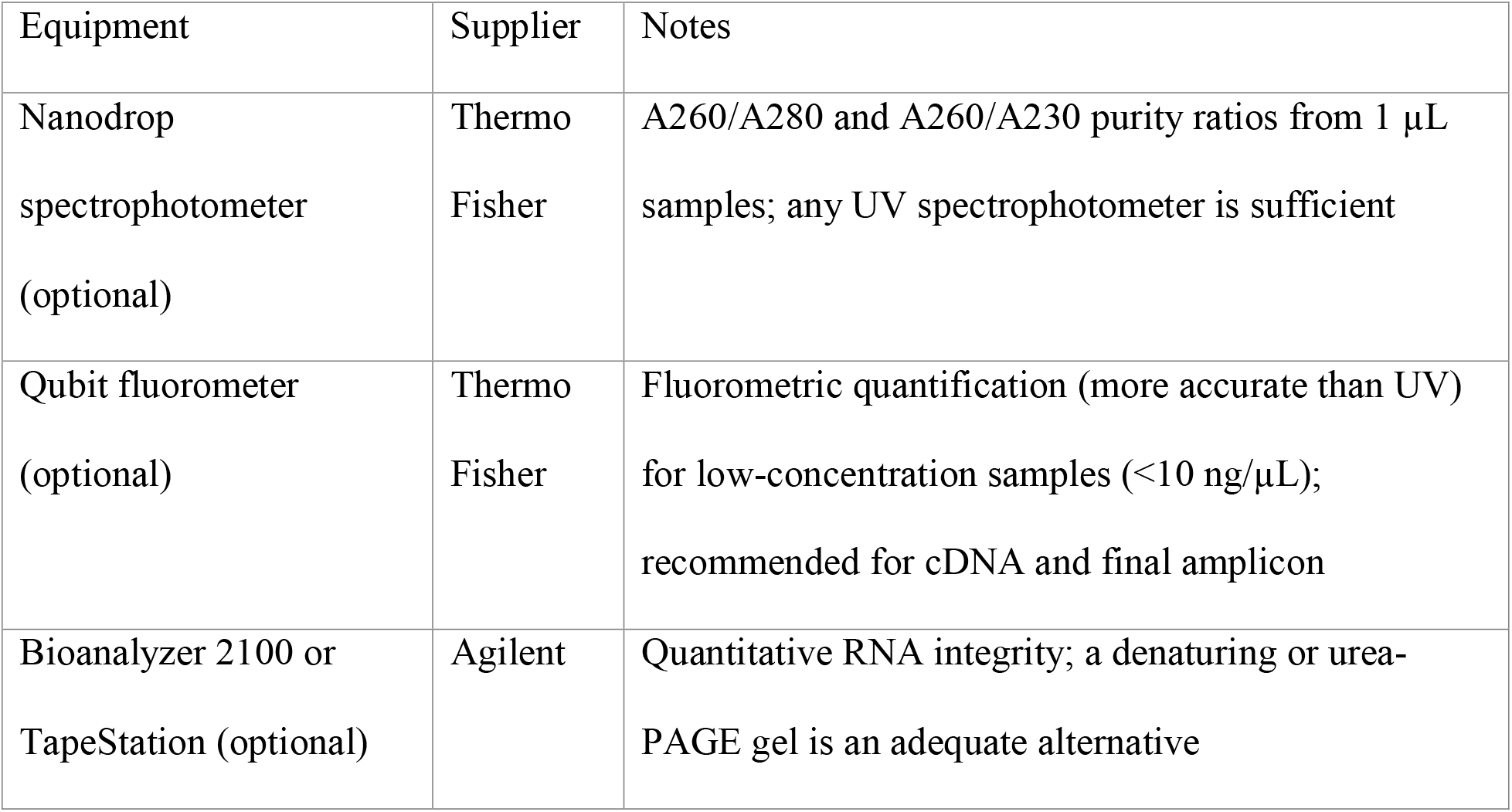

### Software and analysis tools

#### cmuts web server

Browser-based interface wrapping the full FAST-MaP pipeline. Available at https://huggingface.co/spaces/daslab-stanford/cmuts. No local software installation required; all dependencies are handled server-side.

#### ChimeraX (UCSF)

Molecular visualization software used in this work for projecting per-nucleotide reactivity values onto three-dimensional RNA structures (**Fig. 4**). Available at https://www.rbvi.ucsf.edu/chimerax/. Free for academic use.

**Figure 4.**
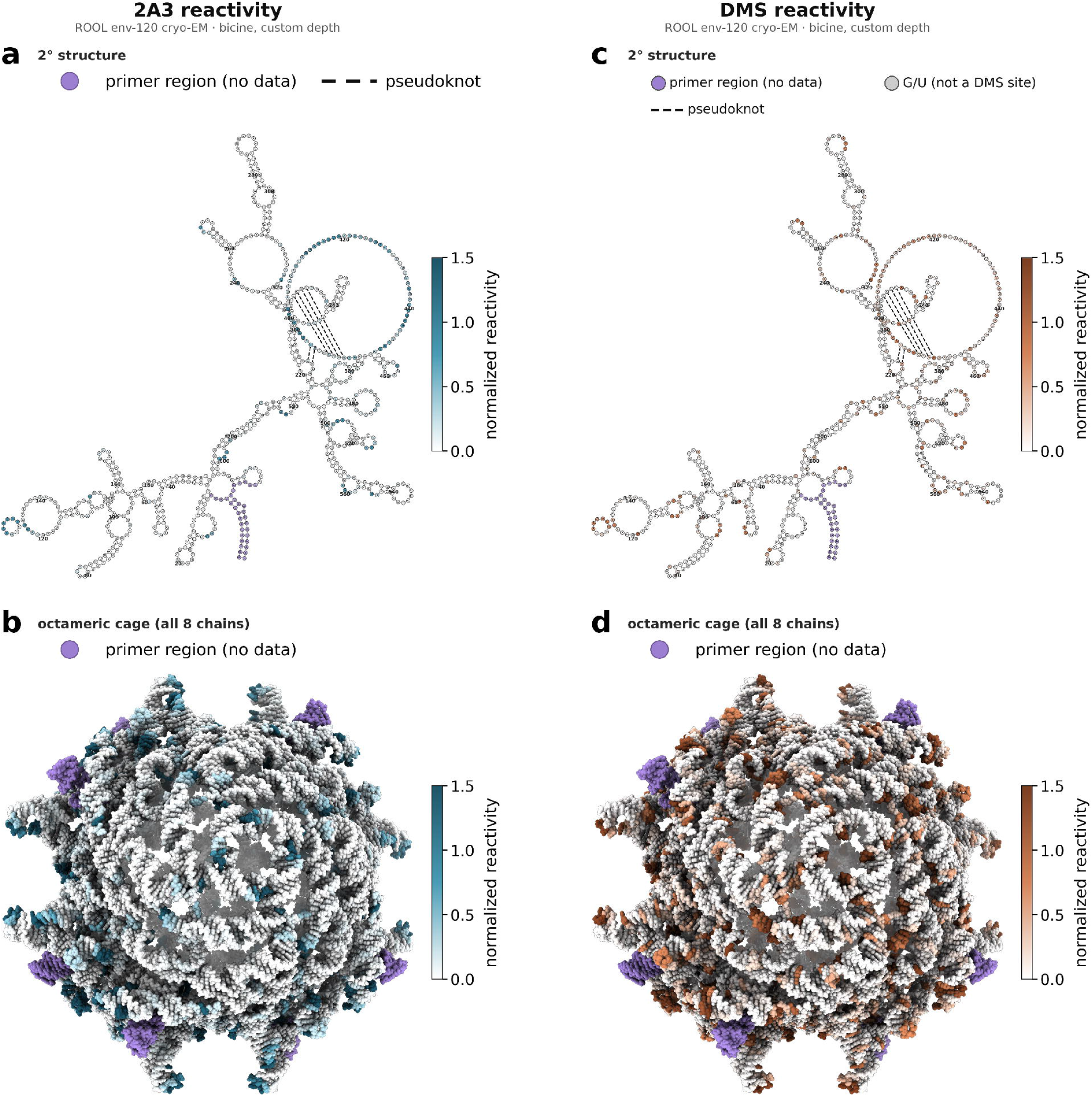
FAST-MaP reactivity mapped onto the ROOL env-120 secondary structure and 3D structure derived from cryo-EM. (a) 2A3 reactivity (teal; bicine, custom depth) mapped onto the ROOL env-120 secondary structure derived from the cryo-EM model (PDB 9MDS, chain A; canonical base pairs including wobble and three pseudoknot layers, verified with RNApdbee/RNApolis) and drawn with forna; pseudoknot pairs are shown as dashed lines. (b) 2A3 reactivity mapped onto all eight chains of the ROOL env-120 octameric cage (PDB 9MDS; structure reproduced from ref. 10 (Kretsch et al.) with permission). (c,d) The same secondary structure and octameric cage colored by DMS reactivity (coral; bicine, custom depth); for DMS the non-reactive G and U positions are shown in grey. All panels use the same reactivity color scale (0–1.5), shown per panel - teal for 2A3 (a,b) and coral for DMS (c,d); pseudoknots (dashed) and non-reactive G/U positions (grey) are marked only on the secondary-structure panels (a,c). Coloring the full assembly rather than a single chain shows the inter-chain quaternary environment and is presented as a structural hypothesis tested by the solution reactivity; intensely colored nucleotides correspond to solvent-accessible, single-stranded regions with high reactivity. Reactivity is reported for the internal region (reference positions 1–615, transcript nt 21–635); the primer-binding regions lie outside this window and are shown in lavender (not analyzed). Throughout, nucleotides use the reactivity numbering 1–615 (the `cmuts` output numbering; transcript positions 21–635 and PDB 9MDS residues 21–635 - add 20 to convert a 1–615 position to its PDB residue number). Quantitative comparison of reactivity at paired vs unpaired nucleotides (violin plots), ROC discrimination, and nucleotide-resolved DMS reactivity are provided as **Supplementary Fig. 4**.

#### PyMOL (Schrödinger)

Alternative molecular visualization software suitable for structure-based reactivity mapping. Available at https://pymol.org. Free educational licenses are available.

#### Sequencing service

Any commercial primer-less sequencing service that accepts purified dsDNA amplicons, performs library preparation internally, and returns FASTQ files. In this work, Plasmidsaurus (Nanopore-based) was used; the protocol is compatible with other sequencing providers meeting these criteria.

### Reagent Setup

#### 2× Marathon Buffer

Prepare fresh before each experiment. Combine the following components in order in a nuclease-free microcentrifuge tube (1 mL total volume):

| Component | Stock conc. | Volume (μL) | 2× conc. |
| --- | --- | --- | --- |
| Tris-HCl, pH 9.0 | 1 M | 100 | 100 mM |
| KCl | 2 M | 200 | 400 mM |
| DTT | 100 mM | 100 | 10 mM |
| Glycerol (sterile) | 80% (v/v) | 500 | 40% (v/v) |
| MnCl<sub>2</sub> | 100 mM | 40 | 4 mM |
| Nuclease-free water | – | 60 | – |
| Total |  | 1,000 |  |

**CRITICAL** Add MnCl_2_ last to avoid precipitation.The 2× Marathon buffer must be assembled fresh before use, as its DTT is not stable on storage. In the RT reaction, 20 µL of 2× buffer is diluted to 1× (50 mM Tris-HCl pH 9.0, 200 mM KCl, 5 mM DTT, 20% glycerol, 2 mM MnCl_2_).

#### Na-HEPES, 0.1 M, pH 8.0 (working stock)

Dilute 1 M Na-HEPES (pH 8.0) 1:10 in nuclease-free water. Store at room temperature, or at −20 °C for longer-term storage. Filter-sterilize and store at room temperature for up to 1 month.

#### NaOH, 0.4 M

Dilute 10 M NaOH 1:25 in nuclease-free water. Prepare fresh before each alkaline hydrolysis step; NaOH solutions absorb atmospheric CO_2_ over time, which reduces their effective concentration.

#### Acid-quench solution^22^

Combine 2 volumes of 5 M NaCl, 2 volumes of 2 M HCl, and 3 volumes of 3 M sodium acetate (NaOAc, pH 5.2). The acid-quench solution contains no labile components and is stable at room temperature; it does not need to be prepared fresh. Each reaction requires 24 µL of stop mix (approximately 7 µL of 5 M NaCl, 7 µL of 2 M HCl, and 10 µL of 3 M NaOAc); scale up proportionally with 10% excess for the number of reactions in the experiment.

#### Ethanol, 70% (v/v)

Combine 7 volumes of 100% molecular-biology-grade ethanol with 3 volumes of nuclease-free water. Prepare fresh or store at −20°C for up to 1 month.

#### DMS working stock

Prepare immediately before use in a chemical fume hood. For this protocol (659-nt RNA), combine 1 µL of 100% DMS with 24 µL of 100% ethanol to yield a 4% (v/v) DMS stock (1% final in the 20 µL reaction). For shorter RNAs, a higher concentration (e.g. 12% stock for 3% final) may be appropriate; see Experimental Design.

**CAUTION** DMS is toxic, volatile, and a suspected carcinogen. Handle only in a chemical fume hood with appropriate personal protective equipment (gloves, lab coat, safety glasses). Dispose of DMS waste according to institutional chemical safety guidelines.

### Equipment Setup

#### Thermocycler programs

The following programs are used throughout the protocol. Program them before starting to minimize delays during experiments.

| Program name | Steps | Used at |
| --- | --- | --- |
| Template PCR | 98°C 30 s; 35× (98°C 10 s, 58°C 20 s, 72°C 30 s); 72°C 5 min; 4°C hold | Stage 1, Step 2 |
| RNA folding | 90°C 3 min; ramp to 25°C over 10 min; 50°C 30 min; ramp to 25°C over 10 min | Stage 4, Step 9 |
| SSII RT (2A3) | 42°C 3 h; 70°C 15 min; 4°C hold | Stage 7, Step 17 |
| Marathon RT (DMS) | 42°C 3 h; 95°C 1 min; 4°C hold | Stage 7, Step 19 |
| Alkaline hydrolysis | 90°C 3 min; 25°C hold | Stage 8, Step 20 |
| cDNA PCR | 98°C 30 s; 23× (98°C 5 s, 55°C 5 s, 72°C 15 s); 72°C 10 min; 4°C hold | Stage 9, Step 23 |

#### E-gel or agarose gel setup

Use 2% agarose with SYBR Safe or equivalent stain. For ROOL env-120, the expected template PCR amplicon (Step 2) is 679 bp; the cDNA PCR amplicon (Step 24) corresponds to the RNA length (659 bp for ROOL env-120), as the T7 promoter is not present in the RNA template. E-gels (Thermo Fisher) provide a convenient pre-cast alternative; run according to the manufacturer’s instructions.

#### Nanodrop / UV spectrophotometer

Blank with the appropriate buffer (nuclease-free water for RNA after ethanol precipitation; Buffer EB for DNA after column purification). For concentrated RNA samples (>1,000 ng/µL), dilute 1:20 before measurement to ensure readings fall within the instrument’s linear range.

##### Box 1

**Choosing a Chemical Probe (DMS vs 2A3)**

FAST-MaP uses two orthogonal chemical probes that report on different structural features. Users may employ both probes for maximum information or select one based on their experimental goals and resource constraints.

**DMS (dimethyl sulfate)** methylates the Watson-Crick face of unpaired adenosine (N1-A) and cytidine (N3-C), directly reporting on base-pairing status. DMS is the recommended starting probe for most applications because it yields higher signal-to-noise at lower read depths (SNR ≈ 3.6–4.5 at pilot sequencing of ∼10,000 reads for ROOL env-120), produces a mutation-based readout via Marathon RT that generates predominantly full-length cDNA, and is the most widely used chemical probe in the RNA structure community, facilitating comparison with published datasets. DMS data are sufficient to identify paired and unpaired A and C residues with high confidence but provide no direct information on G and U base-pairing.

**2A3 (2-aminopyridine-3-carboxylic acid imidazolide)** acylates the 2’-OH of flexible nucleotides at all four bases, reporting on backbone dynamics rather than base-pairing face accessibility. 2A3 is recommended as a complementary probe when users need information at G and U positions, want to detect backbone rigidity at positions that are unpaired but structurally constrained (e.g., stacked single-stranded regions, tertiary contacts), or seek cross-checked structural assignments at A and C positions where both DMS and 2A3 probes report pairing signals. 2A3 requires higher read depths for robust signal (SNR ≈ 1.2–2.5 at pilot; SNR 14.0– 17.0 at ∼416,000–442,000 reads for ROOL env-120), and the deletion-based readout via SuperScript II with Mn^2+^ produces a higher fraction of truncated cDNA, which may reduce PCR amplicon yield.

**Recommendation:** For initial experiments on RNAs above 200 nucleotides, begin FAST-MaP with DMS alone at pilot sequencing to confirm that the protocol is working and to obtain base-pairing information at A and C positions. Add 2A3 with custom sequencing to obtain complementary backbone flexibility data at all four nucleotides. For comprehensive structural characterization, use both probes, as agreement between probes at single-stranded regions strengthens confidence in structural assignments while divergence reveals structural nuance (e.g., stacked but unpaired positions). For RNAs with lengths of 100-200 nts, begin FAST-MaP instead with 2A3 at higher probe concentrations, which provides information at all four nucleotides; then, add custom sequencing and DMS profiling, if desired

##### Box 2

**Ethanol Precipitation Protocol**

The following ethanol precipitation protocol is used at three points in the procedure (Steps 7, 15, and 21). Perform all centrifugation steps at maximum speed in a refrigerated microcentrifuge at 4°C; here, maximum speed denotes the microcentrifuge’s top setting (≥16,000 × g; 21,300 × g on the Eppendorf 5425 used here).

1. Add 1/10 volume of 3 M sodium acetate (NaOAc, pH 5.2), 1 µL of GlycoBlue coprecipitant, and then 2.5 volumes of ice-cold 100% ethanol. Mix thoroughly by vortexing.
2. Incubate at −20°C for at least 30 min.
3. Centrifuge at maximum speed (≥16,000 × g) at 4°C for 30 min. Carefully aspirate the supernatant without disturbing the blue pellet.
4. Wash the pellet twice with 500 µL of ice-cold 70% ethanol, centrifuging at maximum speed for 5 min between washes.
5. Air-dry the pellet for 5–10 min at room temperature. Do not over-dry, as this can make the pellet difficult to resuspend.
6. Resuspend in the volume of nuclease-free water specified in the relevant step.

**Optional QC:** After resuspension, measure concentration by UV absorbance (A260) on any UV spectrophotometer or plate reader. Note that the GlycoBlue co-precipitant absorbs near 260 nm and can inflate UV readings, particularly at low concentrations; when GlycoBlue is used, quantify by a fluorometric assay (e.g., Qubit) for accuracy. If a Nanodrop is available, measure 1 µL directly. Expected A260/A280 for RNA: >2.0; for DNA: ∼1.8.

##### Box 3

**Calculating IVT Reactions**

Determine the number of IVT reactions needed based on the target RNA yield. Each 20 µL MEGAscript T7 reaction typically yields 50–100 µg of RNA, depending on the template and transcript length. To calculate the number of reactions:

1. Estimate the total RNA mass required for all downstream modification reactions (number of conditions × RNA mass per condition).
2. Divide the total required mass by the expected yield per reaction (∼50 µg as a conservative estimate).
3. Round up and add one extra reaction to account for losses during purification and precipitation.

For ROOL env-120 (659 nt), a minimum of 3 reactions is theoretically sufficient (see Box 4); however, 4 reactions are recommended to provide ∼20% excess that compensates for losses during purification and precipitation. In this protocol, we used 4 reactions for the full 8-modification-tube workflow.

##### Box 4

**Worked example calculations (IVT yield, molarity, and resuspension)**

*The following worked example uses ROOL env-120 (659 nt) to illustrate the key calculations. Users should substitute their own RNA length and experimental yields*.

1. **Molecular weight estimation** The average molecular weight of a ribonucleotide is approximately 340 g/mol. For an RNA of length *N* nucleotides: MW = *N* × 340 g/mol For ROOL env-120: MW = 659 × 340 ≈ 224,060 g/mol
2. **Target mass concentration from molar concentration** To convert a target molar concentration (*C*, in µM) to a mass concentration (µg/µL): Mass conc. (µg/µL) = *C* (µM) × MW (g/mol) × 10^−6^ For ROOL env-120 at 34 µM: 34 × 224,060 × 10^−6^ = 7.62 µg/µL ≈ 7.6 µg/µL
3. **Number of IVT reactions needed** Determine the total RNA mass required for all downstream reactions. For FAST-MaP with two buffer conditions (8 modification reactions total, each using 2.35 µL of 34 µM RNA, i.e., 80 pmol per reaction): Total RNA needed = 8 reactions × 2.35 µL × 7.6 µg/µL ≈ 143 µg Using a benchmark yield of ∼57 µg per MEGAscript reaction for a ∼659-nt transcript: Number of reactions = Total RNA needed / Yield per reaction = 143 / 57 = 2.5 → minimum of 3 reactions; recommended 4 reactions to provide ∼20% excess for losses during purification and precipitation. *Note: Include ∼20% excess to account for losses during purification and precipitation steps. Ethanol precipitation recovery is typically 70–90%; factor this into planning if working with limited FAST-MaP requires no special regulatory a*.
4. **Resuspension volume after ethanol precipitation** After measuring the total RNA yield from all IVT reactions (by UV spectrophotometry), calculate the resuspension volume to achieve the target molar concentration: Resuspension vol. (µL) = Total RNA mass (µg) / Target conc. (µg/µL) For ROOL env-120 with a total yield of 227 µg and a target of 7.6 µg/µL (34 µM): Resuspension vol. = 227 / 7.6 = 29.9 µL *TIP: Resuspend in slightly less than the calculated volume, verify the concentration by UV spectrophotometry (dilute 1:20), and add nuclease-free water to reach the target. This avoids overshooting the target concentration, which cannot be corrected without a further precipitation step*.

## PROCEDURE

### Stage 1: PCR amplification of DNA template

#### TIMING ∼10 min setup; ∼1 h thermocycler; ∼30 min gel QC and column purification

Design a forward primer containing the T7 promoter sequence and two initiating guanosine residues (T7GG_F) upstream of the RNA sequence of interest, and a gene-specific reverse primer complementary to the 3′ end of the target RNA. Order a synthetic gene fragment

(e.g. from IDT) encoding the full RNA sequence, including the sequences the primers anneal to (optionally with dedicated 5′/3′ buffer regions). For analysis, the reference FASTA is the full amplicon including these primer-binding regions; cmuts masks the primer positions when their lengths are specified (see Step 27).

1. Prepare a 25 µL PCR reaction using Q5 High-Fidelity 2X Master Mix containing 10 ng of gene fragment template DNA, 0.5 µM each of forward (T7GG_F) and reverse primers, and nuclease-free water to volume.
2. Run the following thermocycling program: initial denaturation at 98°C for 30 s; 35 cycles of 98°C for 10 s, 58°C for 20 s, and 72°C for 30 s; final extension at 72°C for 5 min; hold at 4°C. **NOTE** The annealing temperature of 58°C used here is specific to the T7GG_F/reverse primer pair for ROOL env-120. When adapting this protocol to other RNA targets, the annealing temperature should be recalculated for each new primer pair using an online tool such as the NEB Tm Calculator (https://tmcalculator.neb.com).
3. Load PCR product on a 2% agarose gel (E-gel, SYBR Safe, or equivalent) to verify a single band at the expected full amplicon size (e.g. 679 bp for ROOL env-120, comprising the 659-nt RNA sequence plus the 20-nt T7 promoter and initiating GG dinucleotide). Column-purify the PCR product using a QIAquick PCR Purification Kit: wash twice with Buffer PE, perform a dry spin to remove residual ethanol, and elute in 30 µL of Buffer EB.

**CRITICAL STEP** DNA template purity directly impacts RNA yield and quality in the downstream in vitro transcription reaction. If the A260/230 ratio is below 2.0, repeat the column purification with an additional Buffer PE wash.

**Optional QC:** Measure DNA concentration by UV absorbance (A260). If a Nanodrop is available, expect A260/A280 ≈ 1.8 and A260/A230 > 2.0.

### Stage 2: In vitro transcription (IVT)

#### TIMING ∼30 min setup; 3–4 h incubation

**4.** Assemble the MEGAscript T7 transcription reaction at room temperature to prevent precipitation of the spermidine component. For each 20 µL reaction, combine: 8 µL of a premixed NTP stock containing ATP, CTP, GTP, and UTP at 25 mM each (100 mM total NTP), giving a final concentration of 10 mM each nucleotide (40 mM total), 2 µL of 10× reaction buffer, template DNA at 0.2 µg/µL, 2 µL of enzyme mix, and nuclease-free water to 20 µL. Scale the number of reactions according to **Box 3**.
**5.** Incubate the assembled reactions at 37°C for 3–4 h. Add 1 µL of TURBO DNase to each reaction to digest the DNA template and incubate at 37°C for 15 min.

## TROUBLESHOOTING

**6.** Purify the RNA using a Zymo RNA Clean & Concentrator-25 column. Dilute each reaction to 50 µL with DNase/RNase-free water, add 100 µL of RNA Binding Buffer (2× volume), add 150 µL of 100% ethanol (equal to the combined volume), mix by pipetting, load onto the column, wash according to the manufacturer’s protocol, and elute in 25 µL of DNase/RNase-free water.
**CRITICAL STEP** RNA integrity is essential for obtaining full-length cDNA products during reverse transcription. Degraded RNA will produce truncated cDNA and reduce the signal-to-noise ratio of the final reactivity profile. Do not proceed to chemical modification if the RNA appears degraded.

**Optional QC:** Verify RNA integrity on a denaturing agarose or urea-PAGE gel. A single band at the expected size indicates successful transcription. If a Bioanalyzer or TapeStation is available, RNA integrity number (RIN) >9 is expected. Measure RNA concentration by UV absorbance; expect A260/A280 > 2.0 and A260/A230 > 2.0.

**PAUSE POINT** Purified RNA can be stored at −80°C for several weeks. Avoid repeated freeze-thaw cycles.

### Stage 3: Ethanol precipitation to concentrate RNA

#### TIMING ∼15 min setup; ∼2 h incubation and centrifugation

This step concentrates the RNA to the molar concentration required for the folding reaction. The target concentration depends on the RNA length and molecular weight.

**7.** Combine all IVT eluates into a single microcentrifuge tube and ethanol-precipitate following **Box 2**. Resuspend the pellet in the volume of nuclease-free water calculated to achieve the target molar concentration (34 µM for ROOL env-120; see **Box 3** for determining the required yield). **CRITICAL STEP** Concentrating to a high target (34 µM here) is needed only to fold several conditions in parallel from one preparation. For routine single-RNA probing at ≤1 µM, skip this concentration step and use the RNA directly from the spin-column (e.g. Zymo) eluate, scaling the IVT down (often a single reaction).

**PAUSE POINT** Concentrated RNA can be stored at −80°C for several weeks. Avoid repeated freeze-thaw cycles.

### TROUBLESHOOTING

### Stage 4: RNA folding

#### TIMING ∼15 min setup; ∼1 h thermocycler program; ∼15 min aliquoting

The RNA is thermally denatured and refolded in the presence of Mg^2^+ to achieve a defined secondary and tertiary structure before chemical modification. This protocol was demonstrated using two buffer conditions in parallel (Na-Bicine/KOAc at pH 8.5 and Na-HEPES at pH 8.0) as an optional test of structure preservation across conditions; both yielded well-correlated reactivity profiles (Pearson *r* = 0.895 for 2A3; *r* = 0.933 for DMS), and a single buffer condition may be sufficient for most applications (see Experimental Design for considerations on buffer selection).

**8.** Prepare the folding master mix according to the tables below. For each buffer condition, prepare four reactions (one 2A3-modified, one 2A3-control, one DMS-modified, one DMS-control). Final concentrations listed are for the 20 µL modification reaction (after subsequent addition of 2 µL MgCl_2_ and 5 µL chemical probe).

**Table:** Na-Bicine/KOAc folding reaction setup (per reaction)

**Table: Na-Bicine/KOAc folding reaction setup (per reaction)**
| Component | Stock conc. | Volume (μL) | Final conc. (20 μL) |
| --- | --- | --- | --- |
| RNA (ROOL env-120) | 34 μM | 2.35 | 4.0 μM (80 pmol) |
| Na-Bicine, pH 8.5 | 1 M | 3.00 | 150 mM |
| KOAc | 1 M | 3.00 | 150 mM |
| Nuclease-free water | - | 4.65 | - |
| Subtotal (folding mix) |  | 13.00 |  |
| $\text{MgCl}_2$ (added after refolding) | 100 mM | 2.00 | 10 mM |
| Chemical probe or vehicle (Step 11) | - | 5.00 | - |
| Total reaction volume |  | 20.00 |  |

**Table:** Na-HEPES folding reaction setup (per reaction)

**Table: Na-HEPES folding reaction setup (per reaction)**
| Component | Stock conc. | Volume (μL) | Final conc. (20 μL) |
| --- | --- | --- | --- |
| RNA (ROOL env-120) | 34 μM | 2.35 | 4.0 μM (80 pmol) |
| Na-HEPES, pH 8.0 | 0.1 M | 6.50 | 32.5 mM |
| Nuclease-free water | - | 4.15 | - |
| Subtotal (folding mix) |  | 13.00 |  |
| $\text{MgCl}_2$ (added after refolding) | 100 mM | 2.00 | 10 mM |
| Chemical probe or vehicle (Step 11) | - | 5.00 | - |
| Total reaction volume |  | 20.00 |  |

**9.** Denature the RNA by heating to 90°C for 3 min in a thermocycler. Cool to room temperature over 10 min (ramp down or remove from thermocycler and allow to equilibrate on the bench). Add 2 µL of 100 mM MgCl_2_ to each tube (10 mM in the final 20 µL modification reaction) and mix gently by pipetting. Incubate at 50°C for 30 min to promote tertiary structure formation, then cool to room temperature over 10 min.

**CRITICAL STEP** The folding incubation temperature (50°C) and duration (30 min) may need optimization for your RNA. These conditions were optimized for ROOL env-120 and may not be optimal for all targets. See Experimental Design for guidance on folding condition optimization.

**10.** Aliquot 15 µL of folded RNA into each of the labeled modification reaction tubes (four tubes per buffer condition: 2A3-modified, 2A3-control, DMS-modified, DMS-control).

### Stage 5: Chemical modification with 2A3 and DMS

#### TIMING ∼15 min setup; ∼25 min probe incubation; ∼25 min quenching

Two orthogonal chemical probes are used to interrogate RNA structure. 2A3 acylates the 2′-OH of conformationally flexible (unpaired) nucleotides across all four bases, while DMS methylates the Watson-Crick face of adenosine (N1) and cytidine (N3) at solvent-accessible, unpaired positions. Each probe is applied to one tube of folded RNA, with a matched no-probe control processed in parallel.

**CAUTION** All steps involving DMS must be performed in a chemical fume hood. DMS is volatile, toxic, and a suspected carcinogen. Wear appropriate PPE (nitrile gloves, lab coat, safety glasses). Dispose of DMS-contaminated solid and liquid waste separately according to institutional guidelines.

**11. 2A3 modification:** Prepare a 133.3 mM 2A3 working stock by dissolving 10 mg of 2A3 (MW = 188.19 g/mol) in 133.3 µL of anhydrous DMSO to create a ∼400 mM concentrated stock, then diluting 3-fold (combine 1 volume of 400 mM stock with 2 volumes of anhydrous DMSO; total 3 volumes) to yield the 133.3 mM working stock. To the 2A3-modified tube, add 5 µL of the 133.3 mM 2A3 working stock. To the matched 2A3-control tube, add 5 µL of anhydrous DMSO (vehicle control). Incubate both tubes at room temperature for 20–25 min, mixing by flicking every 5 min to ensure uniform modification.
**12.** Quench the 2A3 reactions by adding 20 µL of 1 M DTT to both the modified and control tubes. Incubate at room temperature for 25 min.

**NOTE** DTT addition is used here as an in-laboratory quench step to scavenge unreacted 2A3 electrophile and halt further acylation prior to RNA recovery, following the original 2A3 reagent paper (ref 3). Users can alternatively quench by immediate dilution and ethanol precipitation; we have found the DTT step to provide more reproducible quenching across replicates.

**13. DMS modification (perform in fume hood):** Prepare fresh 4% (v/v) DMS in ethanol by combining 1 µL of 100% DMS with 24 µL of 100% ethanol (reduced from 12% for this 659-nt RNA; see Experimental Design). Prepare this solution immediately before use. To the DMS-modified tube, add 5 µL of the 4% DMS/ethanol solution. To the matched DMS-control tube, add 5 µL of 100% ethanol (vehicle control). Incubate both tubes at room temperature for 15–20 min in the fume hood, mixing by flicking every 5 min.

**CAUTION** Ensure that all DMS-containing liquids and contaminated consumables (tips, tubes) are disposed of in designated DMS waste containers within the fume hood.

**14.** Quench the DMS reactions by adding 20 µL of 100% β-mercaptoethanol to both the modified and control tubes. Incubate at room temperature for 25 min in the fume hood.

## TROUBLESHOOTING

### Stage 6: Recovery of modified RNA

#### TIMING ∼10 min setup; ∼1.5 h precipitation and centrifugation

**15.** Add nuclease-free water to each tube to bring the total volume to 100 µL. Recover the modified RNA by ethanol precipitation following **Box 2** and resuspend each pellet in 10 µL of nuclease-free water. Use the entire remaining 9 µL as input for the reverse transcription reaction in Step 16.

**NOTE** Ethanol precipitation (Box 2) is recommended for both probes and is the only reliable option for DMS-modified samples: the DMS reaction is strongly acidic, so RNA does not bind a Zymo (silica) column efficiently and is lost during recovery before reverse transcription, and ethanol precipitation also avoids generating additional DMS-contaminated (toxic) waste. For 2A3-modified samples, a Zymo RNA Clean & Concentrator column is an acceptable faster alternative that can improve recovery for longer RNAs.

**PAUSE POINT** Recovered modified RNA can be kept on ice for several hours, or stored at −80°C for up to 1 week. Avoid extended storage of modified RNA; process samples promptly for best results.

**Optional QC:** Remove 1 µL and measure RNA concentration by UV absorbance or fluorometric assay (e.g., Qubit RNA HS). For ROOL env-120, concentrations of 1,000–1,800 ng/µL were obtained. Concentrations substantially below this range may indicate incomplete recovery; use fluorometric quantification (rather than UV absorbance) to confirm, as UV measurements become unreliable at low concentrations due to buffer and salt background signal.

### Stage 7: Reverse transcription

#### TIMING ∼30 min setup (denaturation and annealing); ∼3 h RT incubation; ∼15 min enzyme deactivation

Two different reverse transcriptases are used because the two chemical probes produce structurally informative marks that are best detected by different RT error signatures. 2A3 acylates the 2′-OH, creating a bulky adduct that SuperScript II (in the presence of Mn^2^+) reads out predominantly as deletions, with misincorporations (mutations) occurring at a lower frequency^2,18^. DMS methylates bases (N1-A, N3-C), and Marathon RT, a group II intron-derived enzyme optimized for processivity through modified bases, reads out these modifications predominantly as mutations (misincorporations) rather than truncations^18,19^. Note that both enzymes can produce both mutations and deletions at modified positions. The FAST-MaP pipeline counts both event types and users can configure which events to include during analysis (see Step 29). Using the RT enzyme matched to each probe’s modification chemistry maximizes the signal-to-noise ratio.

**(A) For 2A3-modified samples: reverse transcription with SuperScript II (deletion-based readout).**

The volumes below are 2× the standard SSII reaction to accommodate the larger RNA input volume for this 659-nt construct. The total RT reaction volume is 40 µL.

**Table:** SSII denaturation and primer annealing (per reaction)

**Table: SSII denaturation and primer annealing (per reaction)**
| Component | Stock conc. | Volume (μL) | Notes |
| --- | --- | --- | --- |
| Modified or control RNA | - | 9.0 | Entire remaining sample from Step 15 |
| dNTPs | 10 mM | 4.0 | 1 mM final |
| Nuclease-free water | - | 6.2 |  |
| Subtotal |  | 19.2 | → Denature 90°C 5 min, ice 1 min |
| RT primer<br>(ROOL_120_RP) | 12 μM | 3.0 | 0.9 μM final |
| Subtotal after annealing |  | 22.2 | → Anneal 65°C 10 min, ice 1 min |

**Table:** SSII RT premix (per reaction)

**Table: SSII RT premix (per reaction)**
| Component | Stock conc. | Volume (μL) | Notes |
| --- | --- | --- | --- |
| 5× First-Strand Buffer | 5× | 8.0 | 1× final |
| DTT | 100 mM | 2.0 | 5 mM final |
| MnCl<sub>2</sub> | 100 mM | 2.4 | 6 mM final |
| SuperScript II RT | - | 5.4 |  |
| Total premix | | 17.8 | Add to 22.2 μL RNA/primer mix |
| Total RT reaction |  | 40.0 |  |

**16.** Combine 9 µL of modified or control RNA with 4 µL of 10 mM dNTPs and 6.2 µL of nuclease-free water. Denature at 90°C for 5 min, then transfer immediately to ice for 1 min. Add 3 µL of 12 µM RT primer. Anneal at 65°C for 10 min, then transfer to ice for 1 min.
**17.** Prepare the SSII premix: 8 µL of 5× First-Strand Buffer, 2 µL of 100 mM DTT, 2.4 µL of 100 mM MnCl_2_, and 5.4 µL of SuperScript II enzyme (17.8 µL total). Add the premix to each annealed RNA tube (total reaction volume: 40 µL). Incubate at 42°C for 3 h, then deactivate the enzyme at 70°C for 15 min.

**NOTE** The SSII enzyme volume (5.4 µL) is higher than the manufacturer’s standard usage and was empirically optimized in our laboratory for the 2× scaled 40 µL reaction; we observed improved full-length cDNA yield in the presence of Mn^2+^ at this enzyme loading. Users adapting the protocol to other RNA targets may wish to titrate the SSII volume between 2 and 6 µL per 40 µL reaction.

**CRITICAL STEP** The inclusion of MnCl_2_ in the SSII reaction is essential for promoting deletion-based readout of 2A3 adducts. Without Mn^2+^, SuperScript II will predominantly stall at modified nucleotides rather than bypass them with mutations or deletions, reducing the information content of the sequencing data.

**(B) For DMS-modified samples: reverse transcription with Marathon RT (mutation-based readout).**

The denaturation and primer annealing steps are identical to the SSII protocol above (Step 16), except that no water is added (9 µL RNA + 4 µL dNTPs = 13 µL, then 3 µL primer = 16 µL total before premix addition).

**Table:** Marathon denaturation and primer annealing (per reaction)

**Table: Marathon denaturation and primer annealing (per reaction)**
| Component | Stock conc. | Volume (μL) | Notes |
| --- | --- | --- | --- |
| Modified or control RNA | - | 9.0 | Entire remaining sample from Step 15 |
| dNTPs | 10 mM | 4.0 | 1 mM final |
| Subtotal |  | 13.0 | → Denature 90°C 5 min, ice 1 min |
| RT primer<br>(ROOL_120_RP) | 12 μM | 3.0 | 0.9 μM final |
| Subtotal after annealing |  | 16.0 | → Anneal 65°C 10 min, ice 1 min |

**Table:** Marathon RT premix (per reaction)

**Table: Marathon RT premix (per reaction)**
| Component | Stock conc. | Volume (μL) | Notes |
| --- | --- | --- | --- |
| 2× Marathon Buffer | 2× | 20.0 | 1× final |
| Marathon RT enzyme | - | 4.0 |  |
| Total premix |  | 24.0 | Add to 16 μL RNA/primer mix |
| Total RT reaction |  | 40.0 |  |

**18.** Denature and anneal RNA with dNTPs and RT primer as described in Step 16, but omit the 6.2 µL of nuclease-free water (the Marathon premix volume compensates for this difference).
**19.** Prepare the Marathon premix: 20 µL of 2× Marathon Buffer (Tris-HCl pH 9.0, KCl, DTT, glycerol, MnCl_2_; prepare fresh, see Reagent Setup) and 4 µL of Marathon RT enzyme (24 µL total). Add the premix to each annealed RNA tube (total reaction volume: 40 µL). Incubate at 42°C for 3 h, then deactivate the enzyme at 95°C for 1 min.

**NOTE** The 95°C inactivation temperature is higher than is typical for reverse transcriptases (70– 75°C) and was empirically optimized in our laboratory for cDNA recovery; the brief 1-min exposure at 95°C did not detectably degrade the cDNA in our hands. Users may wish to compare 70°C 15 min and 95°C 1 min inactivation for their RNA target.

### Stage 8: cDNA recovery by alkaline hydrolysis

#### TIMING ∼15 min setup; ∼1 h hydrolysis, neutralization, and precipitation

**20.** To each completed RT reaction, add 13 µL of 100 mM EDTA to chelate divalent cations, followed by 40 µL of 0.4 M NaOH. Incubate at 90°C for 3 min to hydrolyze the RNA template. Allow the tubes to cool to room temperature. Add 24 µL of acid-quench solution (see Reagent Setup) to neutralize the NaOH.
**21.** Ethanol-precipitate the cDNA following **Box 2**. Resuspend the pellet in 8.5 µL of nuclease-free water (7.5 µL is used as PCR input in Step 22; the remainder allows optional QC quantification). NOTE: For long cDNA products, a spin-column concentrator that retains short single-stranded DNA (e.g., Zymo Oligo Clean & Concentrator) may be used instead of ethanol precipitation to recover and concentrate the cDNA.

**PAUSE POINT** Purified cDNA can be stored at −20°C overnight or at −80°C for longer (weeks). cDNA is generally more stable than the source RNA and can be re-quantified before proceeding to PCR if desired. For longer-term storage, transfer the cDNA to DNA LoBind tubes to minimize loss of material to tube walls.

**Recommended QC:** Measure ssDNA concentration by fluorometric assay (e.g., Qubit ssDNA HS) or UV absorbance. Yields will vary depending on the RNA target and modification conditions; for ROOL env-120, concentrations of 85–220 ng/µL were obtained. For samples below 10 ng/µL, fluorometric quantification is strongly recommended, as UV absorbance is unreliable at low concentrations due to buffer and salt background signal; an accurate concentration estimate helps determine the optimal PCR cycle number in the next stage.

## TROUBLESHOOTING

### Stage 9: PCR amplification of cDNA

#### TIMING ∼15 min setup; ∼30 min thermocycler; ∼30 min gel QC and column purification

**22.** Prepare a 25 µL PCR reaction using Phire II Hot Start Master Mix containing 7.5 µL of cDNA, 0.5 µM each of forward primer (ROOL_120_FP, a 20-mer annealing within the 5′ end of the RNA of interest) and reverse primer (ROOL_120_RP, the same reverse primer used for RT and gene fragment amplification in Step 1), and nuclease-free water to volume.

**CRITICAL STEP** For ROOL env-120 the cDNA-amplification primers anneal within the RNA of interest itself (not within dedicated 5′/3′ buffer regions); reactivity values cannot be assigned to nucleotides within the primer-binding footprint because mutations there are overwritten by the primer sequence during PCR. Studies requiring reactivities in the 5′-most or 3′-most positions of the RNA should either include 5′/3′ buffer regions in the gene fragment design (so that primers anneal outside the RNA of interest), or use a primer-ligation strategy in place of nested PCR.

**23.** Run the following thermocycling program: initial denaturation at 98°C for 30 s; 23 cycles of 98°C for 5 s, 55°C for 5 s, and 72°C for 15 s; final extension at 72°C for 10 min; hold at 4°C. **NOTE** The annealing temperature of 55°C is specific to the primer pair and polymerase (Phire II) used here. If using different primers or a different polymerase, recalculate the annealing temperature using the manufacturer’s Tm calculator (e.g., https://tmcalculator.neb.com for NEB polymerases).

**CRITICAL STEP** If the band is faint or absent after gel verification, column-purify the PCR product using a QIAquick column and re-amplify for 10 additional cycles using 5 µL of the purified product as template. This two-step amplification strategy is preferable to simply increasing the cycle number in the initial PCR, as it helps to avoid amplification of non-specific products.

**24.** Load PCR product on a 2% agarose gel to verify a band at the expected size (659 bp for ROOL env-120). If a single clean band is visible, proceed directly to column purification (QIAquick PCR Purification Kit, for single-band products; or MinElute Gel Extraction Kit if gel-extraction is needed). If additional bands are present, gel-extract the band of interest using a MinElute Gel Extraction Kit. Perform two Buffer PE washes, a 2 min dry spin to remove residual ethanol, and incubate the column with 20 µL of Buffer EB for 5 min before eluting.

**PAUSE POINT** Gel-extracted dsDNA amplicon can be stored at −20°C for several weeks before submission to the sequencing service. Re-quantify by fluorometry (e.g., Qubit) or UV absorbance immediately before shipment, as some loss is typical with longer storage.

**Recommended QC:** Measure amplicon concentration by UV absorbance or fluorometric assay for accurate sequencing submission. For ROOL env-120, yields ranged from 11 to 57 ng/µL.

## TROUBLESHOOTING

### Stage 10: Sequencing submission

#### TIMING ∼30 min hands-on

**25.** Dilute the purified dsDNA amplicon to the concentration specified by the sequencing service provider in nuclease-free water. In this work, ROOL env-120 amplicons were submitted at 7 ng/µL in 10 µL; however, input requirements vary between providers, so users should consult their chosen service for current specifications. Submit each sample individually to the sequencing service. No sequencing primers are required; the service ligates sequencing adapters directly to the ends of the dsDNA amplicon.

**NOTE** For ROOL env-120, pilot submissions returned ∼5,700–10,000 reads per sample within approximately 48 h or less, which served as a useful screen to assess sample quality before committing to deeper sequencing. Read counts will vary depending on the provider, submission tier, and amplicon characteristics (see **Provider requirements** in Experimental Design for guidance).

**26.** Download the FASTQ files upon receipt.

**PAUSE POINT** The protocol can be paused here while awaiting sequencing results. Remaining gel-extracted dsDNA can be stored at −20°C.

## TROUBLESHOOTING

### Stage 11: Reactivity profile generation with cmuts

#### TIMING ∼15–30 min hands-on (web server) or ∼1–2 h (command-line installation and analysis)

Upon receipt of FASTQ files from the sequencing service, generate per-nucleotide reactivity profiles using the `cmuts` pipeline^7^. The recommended approach is the `cmuts` web server, which provides a zero-installation browser-based interface that requires no local software (**Fig. 3**). For custom analyses, the `cmuts` command-line pipeline is available as an alternative (see below).

#### Web server analysis (recommended)

**27.** Prepare a reference FASTA file containing the amplicon sequence. The reference determines the coordinates in which the analysis is performed: the first base of the reported reactivity is the first base of the FASTA. For this reason we recommend using the exact sequence which was probed, including the primer binding regions. In that case, specify the primer lengths on the server so `cmuts` masks those positions.

**CRITICAL STEP** A reactivity profile is valid only alongside the reference sequence prepared in this step, so keep the two together in every downstream use.

**28.** Navigate to the `cmuts` web server (**Fig. 3a**). Upload the reference FASTA file and the modified-sample FASTQ file (required). Optionally upload the no-probe control FASTQ file for background subtraction. Unaligned BAM files are accepted in place of FASTQ files.
**29.** Select your sequencing provider in the dropdown, which affects sequence alignment internally. The filtering, counting, subtraction, and normalization settings may remain at their defaults, as they have been tuned specifically for FAST-MaP data. Click “Run Pipeline.”
**30.** On the results page, select the channels that contribute to the final reactivity. Presets reflect the suggested channels for SHAPE (mismatches and deletions) and DMS (mismatches only) chemistry. Inspect the read depth, signal-to-noise ratio, and interactive per-nucleotide reactivity profile (**Fig. 3b**). Download the output files, which are provided in both HDF5 and CSV format for machine learning pipelines and viewing in spreadsheet programs respectively. Cross-check the 2A3 and DMS profiles by examining known structural features such as loops, linkers, and paired regions in the context of the RNA’s known or predicted secondary structure. For ROOL env-120 we derived the secondary structure from the cryo-EM model (PDB 9MDS, chain A) with RNApdbee/RNApolis (canonical pairs including wobble, retaining pseudoknots) and drew it with forna, coloring each nucleotide by reactivity (**Fig. 4a,c**). (Optional) Project the reactivity values onto a three-dimensional structure, if available, by coloring it in ChimeraX (used here) by a per-residue reactivity attribute; for a homo-oligomer, color all chains so that inter-chain interfaces are represented (**Fig. 4b,d**). The cif-overlay tool (github.com/hmblair/cif-overlay) can automate this projection, corroborating the reactivity data with secondary and tertiary structural features (**Fig. 4**).

#### Alternative: Command-line analysis with cmuts

For batch processing of multiple conditions, custom analysis parameters, or integration into existing bioinformatics pipelines, the `cmuts` command-line tools can be used instead of the web server, see the documentation (daslab.stanford.edu/cmuts) for detailed information on options and troubleshooting. The following steps assume `cmuts`, `minimap2`, and `samtools` are installed (see Software and analysis tools).

**(a)** Align FASTQ reads to the reference sequences using `cmuts align`, which wraps `minimap2` for alignment and `samtools` for sorting; `-x` names the sequencing platform: `cmuts align -f references.fasta -x sr -o modified.bam modified.fastq.gz`
**(b)** Run `cmuts hmm` on each sorted alignment to compute per-nucleotide mutation rates, output as an HDF5 file:

~~~
cmuts hmm -f references.fasta -o modified.h5 modified.bam
~~~

**(c)** Run `cmuts sub` to perform background subtraction:

~~~
cmuts sub -o reactivity.h5 modified.h5 control.h5
~~~

**(d)** Run `cmuts norm` to normalize the reactivity:

~~~
cmuts norm -o normalized.h5 reactivity.h5
~~~

**(e)** Verify data quality and visualize the results with `cmuts plot`, which reports read depths, mean reactivity, and per-sample signal-to-noise alongside the reactivity profiles and heatmap:

~~~
cmuts plot normalized.h5 modified.h5 control.h5
~~~

**(f)** (Optional) Project the reactivity values onto a three-dimensional structure with `cif-overlay` (github.com/hmblair/cif-overlay), which colors a CIF structure’s residues per the `cmuts` HDF5 output and renders it in ChimeraX.

## TROUBLESHOOTING

### TIMING

The estimated time for each stage of the FAST-MaP protocol is listed below. Total hands-on time is approximately 24–30 hours spread over 5 working days, plus roughly 6–48 hours for the sequencing service to return data, for an overall time of 1 week from protocol start to analyzed data.

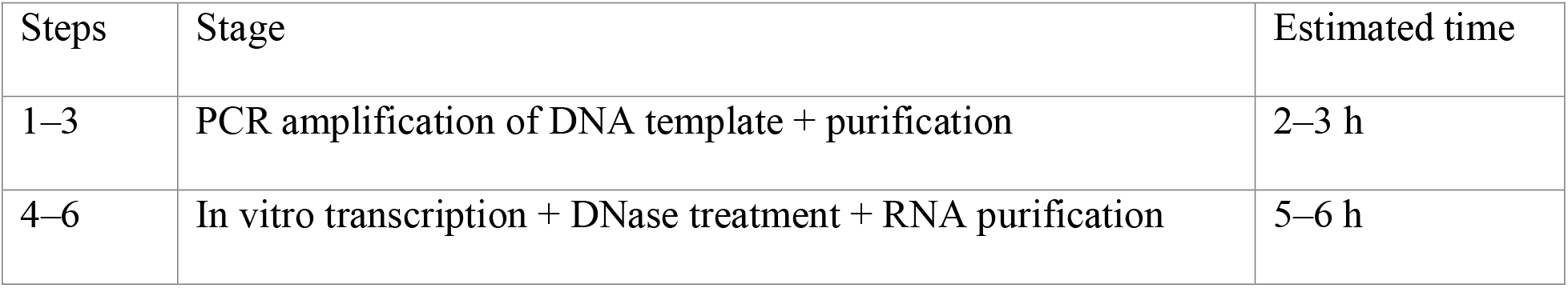
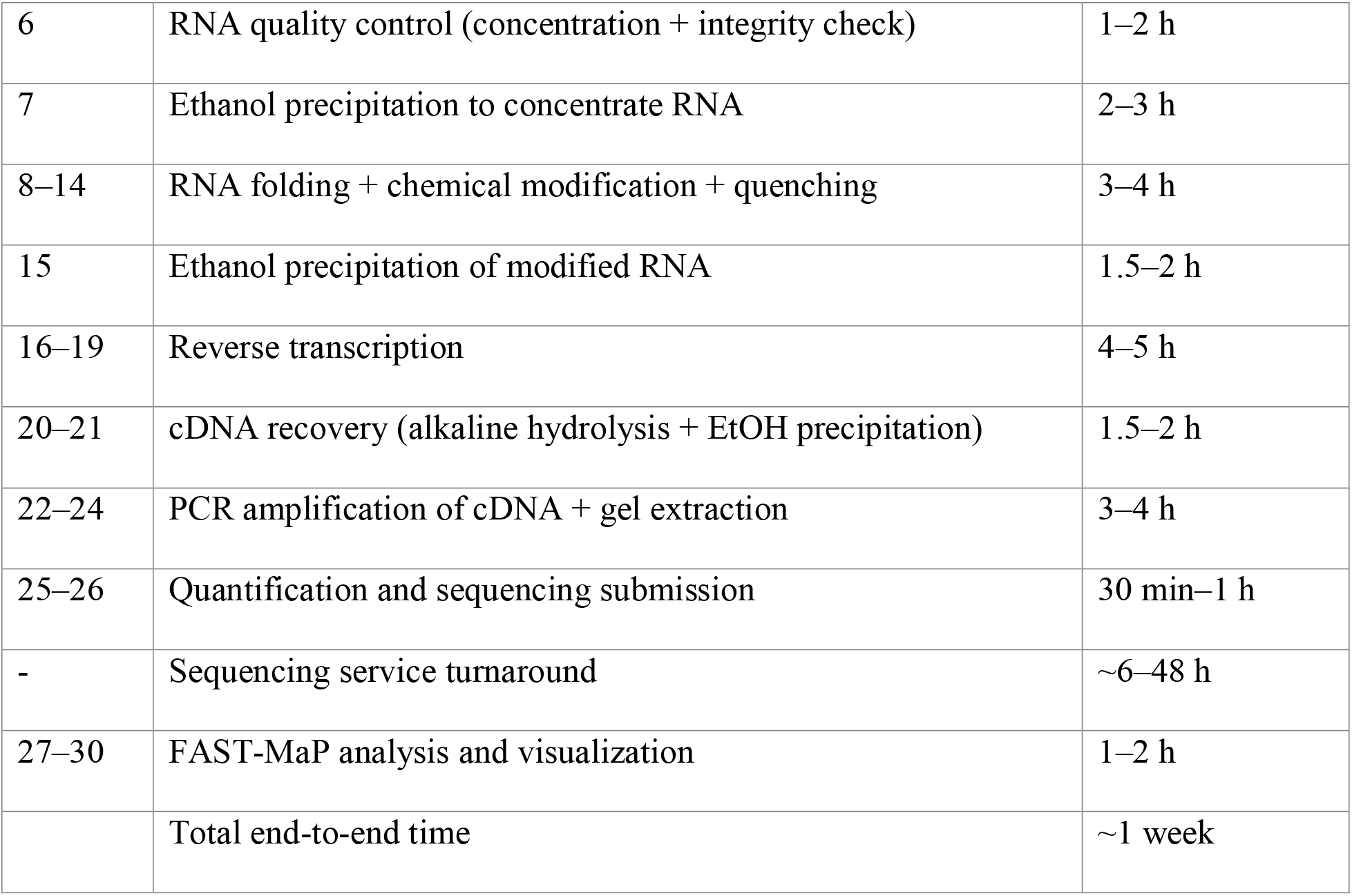

## TROUBLESHOOTING

**Table 2.** Troubleshooting guide for common issues encountered during the FAST-MaP protocol.

| Step | Problem | Possible reason | Solution |
| --- | --- | --- | --- |
| 5 | Low RNA yield after IVT | Suboptimal DNA template quality or insufficient input | Increase number of IVT reactions; verify DNA template purity (A260/280 ~1.8, A260/230 >2.0); extend incubation to 4 h; ensure NTP stocks are not degraded |
| 7 | Poor A260/230 ratio after ethanol | Ethanol or salt carryover in the | Repeat ethanol precipitation with an additional 70% ethanol wash; ensure |
|  | precipitation | RNA pellet | complete air drying (~10 min); avoid over-drying |
| 7 | RNA concentration lower than expected after resuspension | Loss during precipitation or inaccurate UV absorbance measurement | Ensure GlycoBlue is added as co-precipitant; centrifuge at maximum speed for 30 min at 4°C; verify spectrophotometer blank with the correct buffer; cross-check with fluorometry (e.g., Qubit RNA HS) |
| 14 | RNA degradation after chemical modification | Over-modification or prolonged incubation | Reduce probe incubation time (15 min for DMS; 20 min for 2A3); for longer RNAs, reduce 2A3 concentration (133 mM was used for 659-nt ROOL env-120); verify quenching reagents are fresh |
| 21 | Low cDNA yield after reverse transcription | Suboptimal RT conditions or degraded enzyme | Verify RNA input by fluorometry (e.g., Qubit) or UV absorbance; use fresh RT enzyme; ensure Mn<sup>2+</sup> (for SSII) or Marathon buffer is correctly prepared; confirm thermocycler accuracy at 42°C |
| 24 | Faint or absent PCR band after cDNA amplification | Low cDNA input or excessive modification reducing RT | Column-purify the PCR product (QIAquick) and re-amplify for 10 additional cycles using the purified product as template; do not increase the |
|  |  | processivity | initial cycle number above 23 |
| 24 | Multiple bands or smearing on E-gel | Non-specific amplification or primer dimers | Reduce PCR cycle number; verify primer specificity; gel-extract only the band at the expected size; increase annealing temperature by 2–3°C |
| 24 | Apparent ‘protein contamination’ by UV absorbance for DMS/HEPES samples | UV-absorbing artifacts from residual DMS or buffer components | This is a known artifact and does not indicate actual protein contamination. Use fluorometry (e.g., Qubit dsDNA HS) as the primary quantification method |
| 26 | Low read count returned from sequencing service | Insufficient dsDNA input or sample degradation during shipping | Verify concentration by fluorometry (e.g., Qubit) immediately before submission; ensure sample meets the service’s minimum concentration and volume |
| 30 | Low SNR in reactivity profiles | Insufficient read depth or high background in controls | Request higher read depth (custom tier); verify no-probe controls show low background; re-examine RNA quality and modification efficiency |
| 30 | Reactivity profile is shifted | The profile is being read against a sequence other than the reference | Positions are numbered from the first base of the reference provided, so check the profile against that sequence; convert the numbering before you |
|  |  | provided | compare the profile against any other sequence |
| 30 | High background in no-probe control | RNA degradation, residual DNA contamination, or suboptimal RT fidelity | Verify RNA integrity by capillary electrophoresis (e.g., Bioanalyzer) or denaturing gel; confirm complete DNase treatment; re-run RT with fresh enzyme; consider reducing RT incubation temperature |

## ANTICIPATED RESULTS

This section describes expected outcomes at each QC checkpoint. All benchmarks use ROOL env-120 (659 nt; PDB: 9MDS)^10^. Representative data are shown in **Figures 2** and **4** and **Supplemental Data**.

### RNA quality and yield

After in vitro transcription and column purification (Step 2), A260/A280 and A260/A230 should both be >2.0. The integrity check (capillary electrophoresis or denaturing gel) should show a single sharp peak at the expected size with no degradation smear (**Supplementary Fig. 1a**). For ROOL env-120, a RIN of 9.9 was obtained.

After ethanol precipitation (Step 3), resuspend to the target molar concentration (Box 4). For ROOL env-120 at 34 µM, the theoretical mass concentration is ∼7.6 µg/µL; a reading of 8.3 µg/µL was obtained and adjusted to target.

### cDNA and dsDNA library yields

After reverse transcription and cDNA recovery (Steps 7–8), cDNA can optionally be quantified (Qubit ssDNA HS or UV) before PCR (Step 9). DMS samples tend to yield higher cDNA than 2A3 samples, consistent with Marathon RT giving mostly full-length cDNA and SuperScript II giving more shorter fragments due to inefficient mutational bypass (and termination) at 2A3 sites. For ROOL env-120, cDNA ranged 85–220 ng/µL.

After PCR and gel extraction (Step 9), the dsDNA amplicon should appear as a single band at the expected size on a 2% E-gel (659 bp for ROOL env-120). 2A3 samples may give fainter bands; see the CRITICAL STEP at Step 23 for the re-amplification rescue. Capillary electrophoresis traces (**Supplementary Fig. 1b–i**) show a dominant peak at ∼900 bp across all conditions. For ROOL env-120, dsDNA ranged 11–57 ng/µL after gel extraction.

### Sequencing output and read depth

Read depths vary by service and tier. For ROOL env-120, pilot submissions returned ∼5,700– 10,000 reads per sample; a custom order (0.264 Gb, 8 samples) delivered 416,000–520,000 reads per sample; see PCR and sequencing considerations for order-size calculations.

### Reactivity profiles

The normalized reactivity heatmap (**Fig. 2a**) displays per-nucleotide reactivity across all eight conditions at both depths. The key feature is concordance between depths: high- and low-reactivity regions should match, confirming pilot data capture the qualitative profile. Reactivity line profiles (**Fig. 2b**) and the cross-buffer comparison (**Supplementary Fig. 2a**) provide complementary views; the pattern should be consistent between Na-Bicine and HEPES buffers.

### Reproducibility

Cross-buffer reproducibility (**Supplementary Fig. 2a**) provided the primary quality metric for these demonstration samples. For ROOL env-120, Pearson *r* = 0.895 (2A3) and 0.933 (DMS), confirming genuine structural features. Pilot-to-custom reproducibility (**Supplementary Fig. 2b**) was also high (DMS *r* = 0.996; 2A3 *r* = 0.980–0.998), showing pilot depths capture the qualitative pattern despite ∼50-fold fewer reads.

### Signal-to-noise analysis

DMS consistently yields higher SNR than 2A3 at equivalent read depths (**Supplementary Fig. 2c**), as mutation-based readout captures more per-read information than deletion-based readout. At pilot depths, DMS reached SNR 3.6–4.5, sufficient for major structural features; 2A3 gave ≈ 1.2–2.5, approaching the limit of usability to distinguish paired from unpaired nucleotides. At custom depths, both reached excellent SNR (Table 3).

**Table 3:** Sample-level signal-to-noise ratio (SNR) benchmarks for ROOL env-120, as reported by cmuts plot (reads and SNR at Min mapping quality 20 (cmuts CLI default), ubr normalization; DMS = mismatch only, 2A3 = mismatch + deletion)

| Sequencing | Condition | Reads | SNR | Probe | ROC-AUC |
| --- | --- | --- | --- | --- | --- |
| Pilot | Bicine | 5,703 | 1.16 | 2A3 | 0.654 |
| Pilot | HEPES | 9,972 | 2.47 | 2A3 | 0.656 |
| Pilot | Bicine | 9,973 | 4.50 | DMS | 0.790 |
| Pilot | HEPES | 9,987 | 3.61 | DMS | 0.792 |
| Custom | Bicine | 442,026 | 17.03 | 2A3 | 0.725 |
| Custom | HEPES | 416,379 | 13.95 | 2A3 | 0.690 |
| Custom | Bicine | 519,819 | 29.80 | DMS | 0.799 |
| Custom | HEPES | 477,127 | 22.77 | DMS | 0.821 |

Sample-level SNR (**Table 3**; **Supplementary Fig. 2c**) scales roughly with the square root of read count. Per-nucleotide precision varies with base identity, pairing, and depth (**Supplementary Fig. 3**): median reactivity-to-error ratios ranged 2.2–2.8 (DMS, pilot) to 12.8–17.1 (DMS, custom) and 0.7–1.0 (2A3, pilot) to 3.6–6.4 (2A3, custom). Pilot depth gives a qualitative overview; custom depth is needed for precise quantitation.

### A failure mode and its diagnosis (problematic-data example)

As a worked under-powered example, the pilot 2A3 profiles for ROOL env-120 fell to SNR ≈ 1.2–2.5 at ∼5,700–10,000 reads (Bicine 1.16; HEPES 2.47; see **Table 3**), near the usability threshold. The profile keeps the qualitative pattern of the custom data (cross-depth Pearson *r* = 0.980–0.998) but per-nucleotide values fluctuate between replicates. 2A3 (SuperScript II) yields fewer informative events per site than DMS (Marathon RT), so more reads are needed. The fix here was to re-submit the 2A3 samples at a higher tier (see **Read depth**); the DMS data from the same submission were usable at pilot depth. This matches the pattern in **Box 1**: use DMS at pilot depth to confirm the protocol works, then add 2A3 with a custom follow-up order.

### Testing a cryo-EM model with FAST-MaP reactivity

When a 3D structure is available, reactivity can be projected onto it for visualization and comparison (**Fig. 4**; detailed analysis in **Supplementary Note 1**). We mapped FAST-MaP reactivity both onto the secondary structure derived from the cryo-EM model (drawn with forna) and onto the 3D structure formed by all eight chains of the octameric ROOL cage (**Fig. 4a–d**).

For ROOL env-120, high-reactivity nucleotides clustered at flexible regions and low-reactivity ones at helical stems, and ROC analysis confirmed paired/unpaired discrimination (AUC: 2A3 = 0.723; DMS = 0.815; **Supplementary Fig. 4b**). In addition, qualitative inspection reveals protections from 2A3 and DMS reactions in some loops in the secondary structure (**Fig. 4a,c**), which support the formation of loop-loop interactions seen in the ROOL tertiary and quaternary structure (**Fig. 4b,d**).

Together, these results show that established chemical-probing chemistry paired with commercial primer-less sequencing produces high quality reactivity profiles in about one week using standard equipment and widely accessible commercial services. FAST-MaP’s use of `cmuts` pipeline and web server further lowers the barrier via zero-installation analysis.

## Supporting information

FAST-MaP_Supplementary_Information

FAST-MaP_ROOL-env120_Supplementary_Data

## AUTHOR CONTRIBUTIONS

J.V. conceived the FAST-MaP protocol, designed and performed the chemical probing experiments, generated benchmarking data, drove development of the `cmuts` web server, and wrote the manuscript. H.M.B. developed the `cmuts` analysis software, pipeline, and the `cmuts` web server. W.K. provided expertise in chemical mapping methodology and experimental guidance during protocol development. R.D. supervised the project and edited the manuscript.

## ACKNOWLEDGEMENTS

This work was supported by the National Institute of General Medical Sciences (NIGMS), National Institutes of Health (R35 GM122579 to R.D.). R.D. is a Howard Hughes Medical Institute Investigator. The authors used generative AI tools (Anthropic Claude) to assist with drafting and editing the manuscript, preparing figures, and writing data-analysis and visualization code; all AI-assisted content was reviewed and verified by the authors, who take full responsibility for the work.

## COMPETING INTERESTS

J.V., H.M.B., and R.D. filed an invention disclosure with Stanford University covering the FAST-MaP protocol and associated analysis tools. The authors declare no other competing interests.

## DATA AVAILABILITY

Processed per-nucleotide reactivity profiles (with standard error and read counts), the `cmuts` analysis inputs and outputs, the secondary and tertiary structures used for the figures, and the raw pilot sequencing reads for ROOL env-120 are provided as Supplementary Data (`FAST-MaP_ROOL-env120_SupplementaryData.zip`); a complete file listing is given in the Supplementary Information. These data are also provided as Source Data for Figures 2 and 4.

The custom high-depth raw sequencing (FASTQ) is available from the authors on request.

## CODE AVAILABILITY

The `cmuts` analysis pipeline is freely available at https://github.com/DasLab/cmuts under the MIT License. A versioned snapshot of the `cmuts` code corresponding to this protocol is archived on Zenodo (v2.0.0: 10.5281/zenodo.22805311; all versions: 10.5281/zenodo.22805310); the current version is maintained at the GitHub repository above. The `cmuts` web server is accessible at https://huggingface.co/spaces/daslab-stanford/cmuts and requires no installation or account creation.

## SUPPLEMENTARY INFORMATION

The following items accompany this manuscript:

- **Supplementary Figure 1** | Bioanalyzer quality-control traces for ROOL env-120 RNA and gel-extracted dsDNA amplicons (panels a–i).
- **Supplementary Figure 2** | Cross-buffer and cross-depth reproducibility, and signal-to-noise scaling for ROOL env-120.
- **Supplementary Figure 3** | Per-nucleotide precision (reactivity / standard error) characterization, with summary statistics for all four probe-buffer conditions at both sequencing depths.
- **Supplementary Figure 4** | Quantitative agreement between FAST-MaP reactivity and the ROOL env-120 cryo-EM secondary structure: violin plots, ROC, and nucleotide-resolved DMS reactivity.
- **Supplementary Note 1** | Detailed comparison of FAST-MaP reactivity with the ROOL env-120 cryo-EM structure (PDB 9MDS), including DSSR base-pairing assignment commands and the ChimeraX reactivity-mapping command (setattr #1 res bfactor reactivity_value, then color by attribute bfactor). This Note explains the analyses in **Supplementary Figs. 2 and 4**.
- **Supplementary Table 1** | Oligonucleotide primer sequences used in this protocol (T7GG_F, ROOL_120_FP, ROOL_120_RP).

## Notes

https://huggingface.co/spaces/daslab-stanford/cmuts

