## Supplementary material for "FAST-MaP: Chemical Mapping of RNA Structures Using Primer-less Sequencing": FAST-MaP_Supplementary_Information

#### Supplementary Figures

**Supplementary Figure 1 | Bioanalyzer quality-control traces for ROOL env-120 RNA and gel-extracted dsDNA amplicons (panels a–i).**

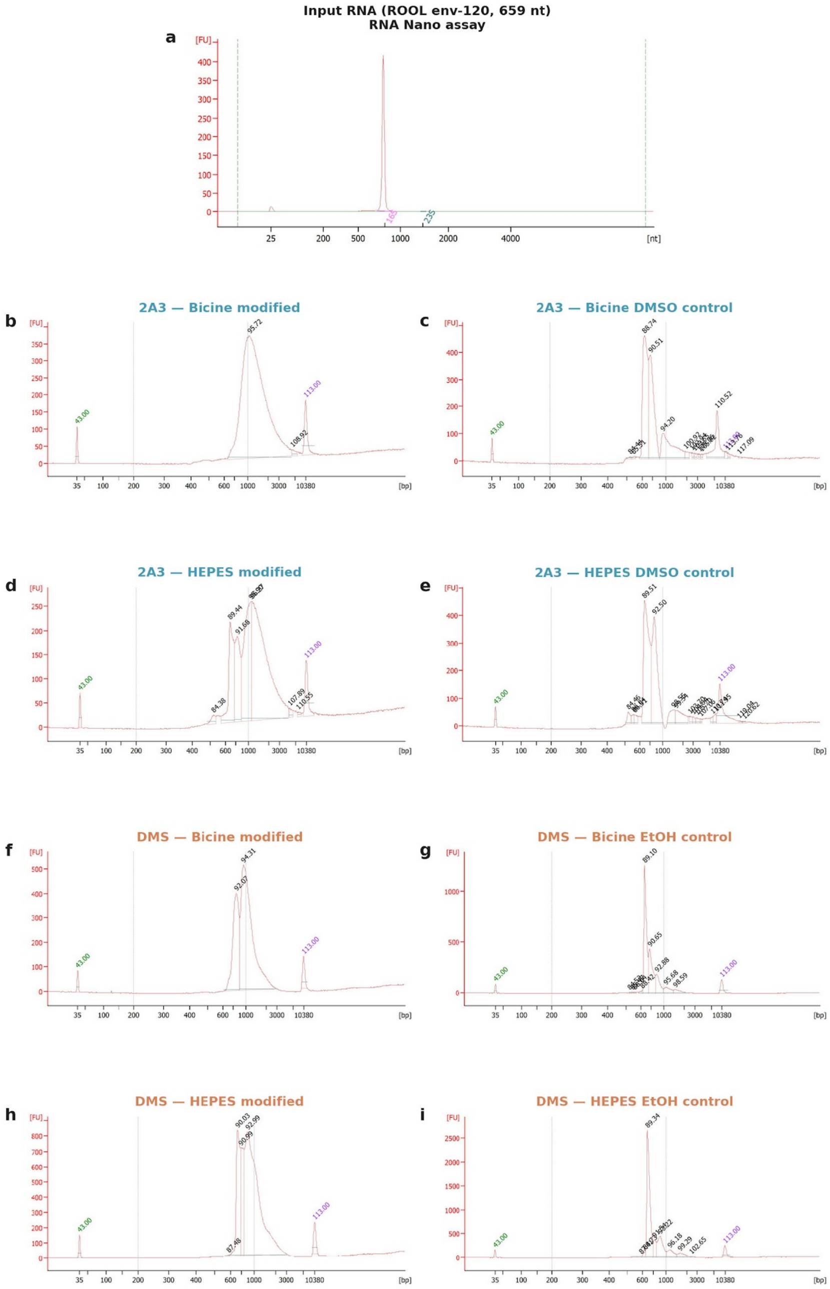


*Related to Figure 2.*

(a) Agilent Bioanalyzer electropherogram (RNA Nano assay) of the input RNA (in vitro-transcribed ROOL env-120, 659 nt), prior to chemical modification. A single dominant peak at the expected migration time confirms RNA integrity. (b–c) 2A3-Bicine: modified sample (b) and DMSO no-probe control (c). (d–e) 2A3-HEPES: modified sample (d) and DMSO no-probe control (e). (f–g) DMS-Bicine: modified sample (f) and EtOH no-probe control (g). (h–i) DMS-HEPES: modified sample (h) and EtOH no-probe control (i). Teal panel titles indicate 2A3 conditions; coral panel titles indicate DMS conditions. All amplicons show a dominant peak at the expected size. DMS-derived amplicons (Marathon RT) typically display a sharper peak, whereas 2A3-derived amplicons (SuperScript II) may show a broader profile reflecting truncation at modification sites during reverse transcription.

**Supplementary Figure 2 | Cross-buffer and cross-depth reproducibility, and signal-to-noise scaling for ROOL env-120.**

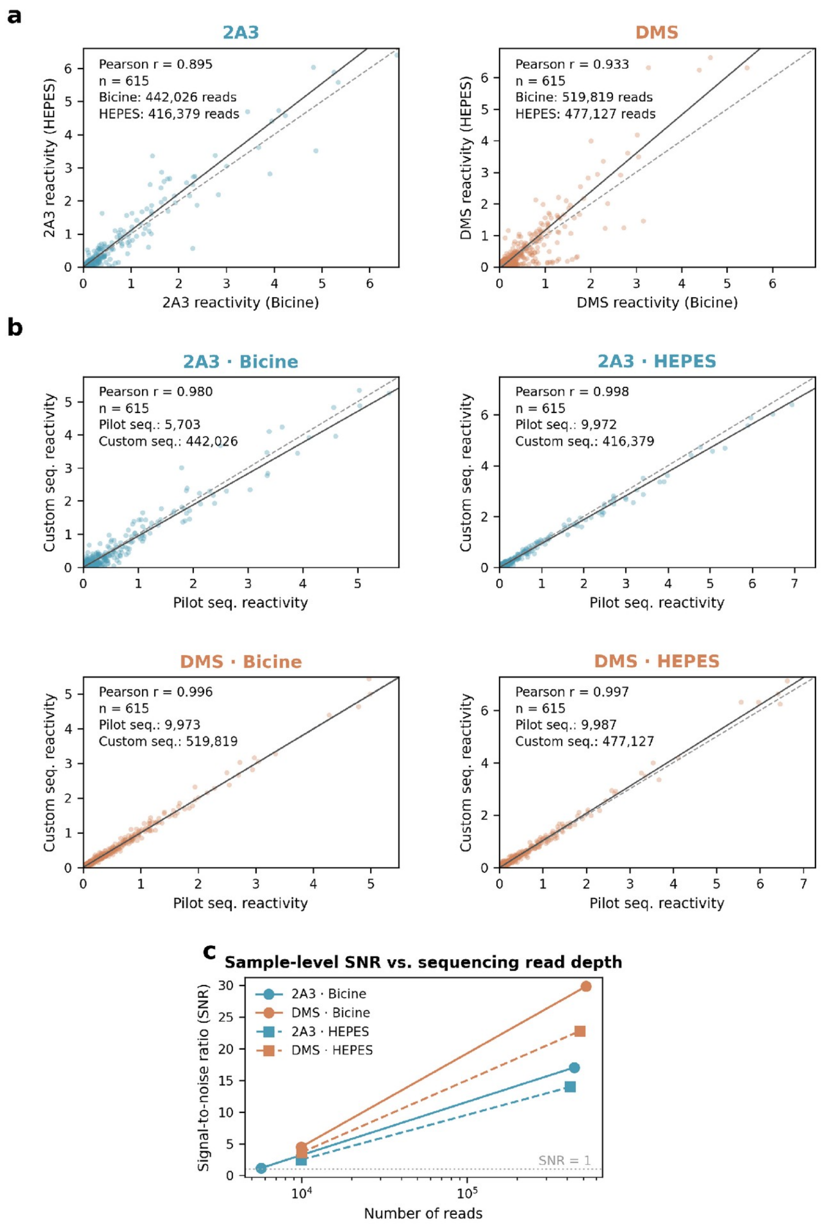


*Related to Figure 2.*

(a) Cross-buffer reproducibility scatter plots comparing Bicine versus HEPES reactivity for 2A3 and DMS at higher sequencing depth (~416,000–520,000 reads/sample). Pearson r values, linear fits, and read counts are shown. (b) Pilot versus custom sequencing reproducibility scatter plots for all four probe-buffer combinations (2×2 grid), demonstrating high concordance even at ~50-fold differences in read depth. (c) Sample-level signal-to-noise ratio (SNR), as reported by the FAST-MaP pipeline, as a function of sequencing read depth for each probe-buffer combination. DMS consistently achieves higher SNR than 2A3 at equivalent depths. The dashed line marks SNR = 1. Note that this is an aggregate sample metric; per-nucleotide precision varies by position (see Supplementary Fig. 3).

**Supplementary Figure 3 | Per-nucleotide precision (reactivity / standard error) characterization, with summary statistics for all four probe-buffer conditions at both sequencing depths.**

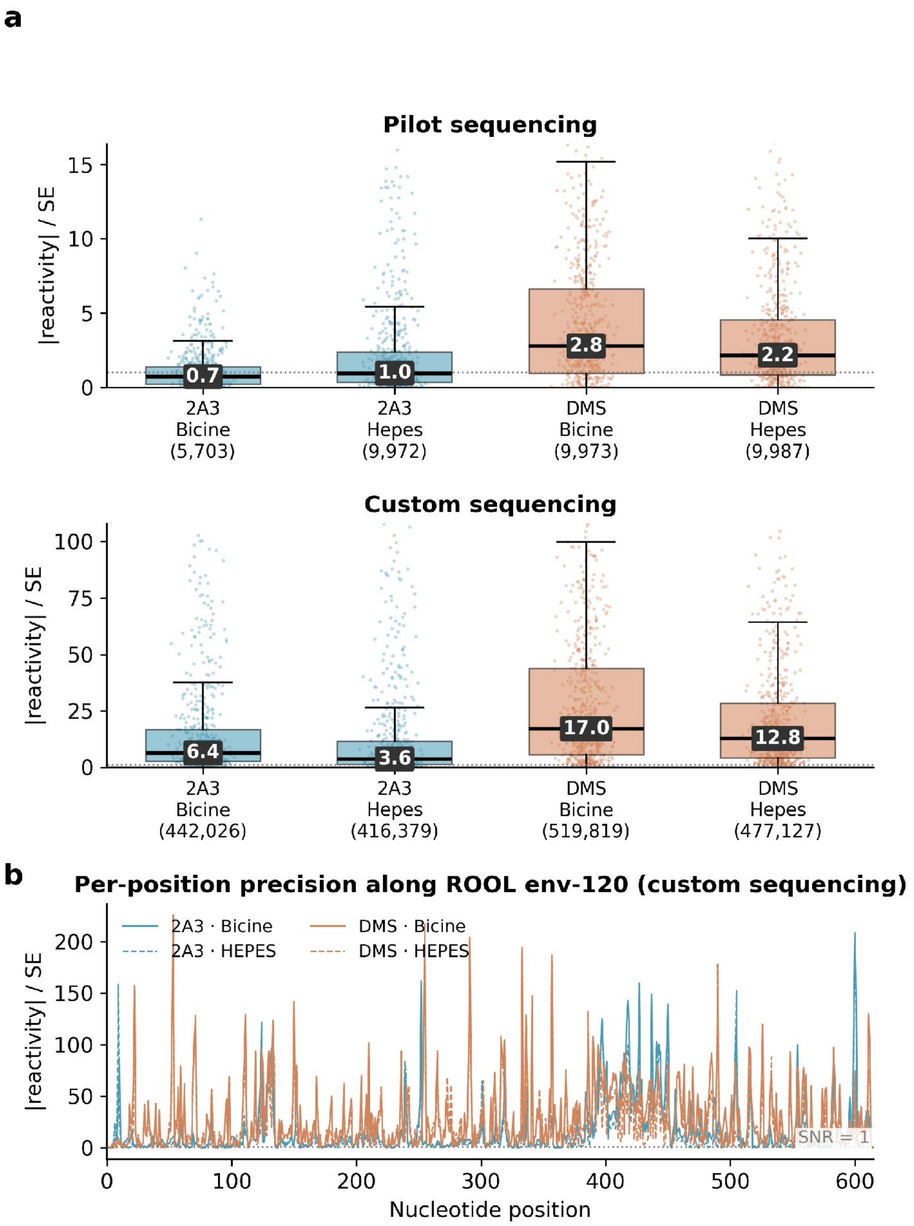


*Related to Figure 2.*

(a) Box-and-whisker plots showing the distribution of per-nucleotide precision, defined as |reactivity| / standard error, for all four probe-buffer conditions. Top row: pilot sequencing (~5,700–10,000 reads), with the y-axis limited to the whisker range (extreme outliers omitted) to avoid compressing the boxes. Bottom row: custom sequencing (~416,000–520,000 reads), showing markedly improved precision. Median values are annotated on each box; read counts are shown below each condition. The dotted line marks a ratio of 1 (signal equals noise). At pilot depth, median per-nucleotide ratios are below 3 for all conditions; at custom depth, DMS conditions achieve median ratios of 13–17, while 2A3 conditions reach 3.6–6.4. (b) Per-position precision trace along the ROOL env-120 reference (615 nt) at custom sequencing depth. The raw per-position ratio is shown (no smoothing applied). DMS (coral) shows consistently higher per-position precision than 2A3 (teal), with Bicine (solid) and HEPES (dashed) buffers showing similar profiles. The dotted line marks SNR = 1. n = 615 nucleotides per condition.

**Supplementary Figure 4 | Quantitative agreement between FAST-MaP reactivity and the ROOL env-120 cryo-EM secondary structure: violin plots, ROC, and nucleotide-resolved DMS reactivity.**

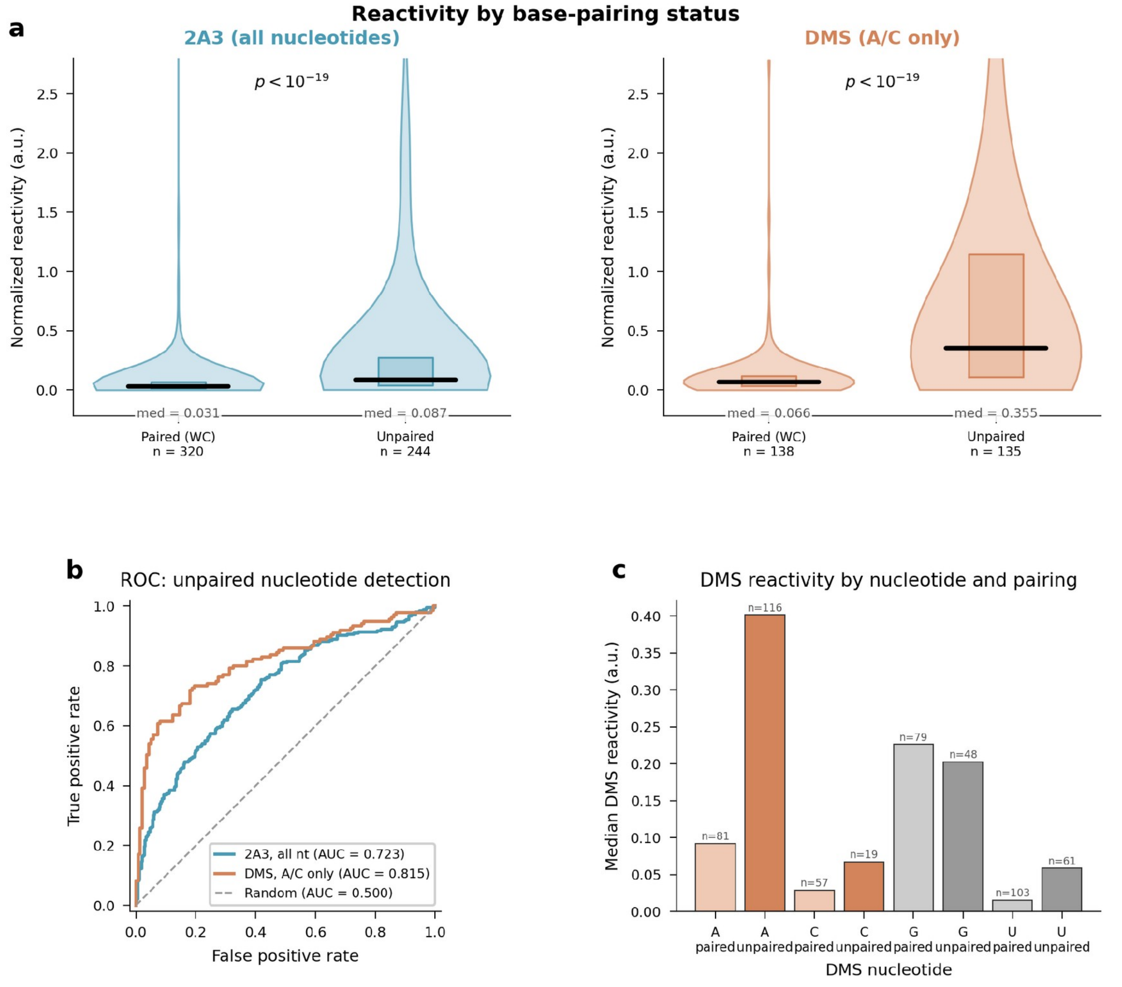


*Related to Figure 4.*

(a) Violin plots comparing normalized reactivity distributions for paired (Watson-Crick base-paired) versus unpaired nucleotides, as classified from the cryo-EM structure (PDB 9MDS). Left: 2A3 reactivity for all nucleotides (teal); right: DMS reactivity restricted to A and C nucleotides (coral). Both probes show significantly higher reactivity at unpaired positions (Mann-Whitney U test, P values indicated). Median values and IQR boxes are overlaid; sample sizes shown below each category. (b) Receiver operating characteristic (ROC) analysis evaluating the ability of 2A3 and DMS reactivity to discriminate unpaired from paired nucleotides. AUC values: 2A3 (all nt) = 0.723; DMS (A/C only) = 0.815. Random classifier (AUC = 0.500) shown as dashed diagonal. (c) Median DMS reactivity by base identity and pairing. A shows the strongest pairing dependence (unpaired ⋙ paired), with C, and to a similar degree U, showing weaker but consistent unpaired-over-paired reactivity, consistent with minor N3-U methylation (ref 21). G shows uniformly elevated reactivity independent of Watson-Crick pairing, consistent with DMS modification at the N7 atom, which is on the Hoogsteen and not Watson-Crick edge of G (ref 23). Because A and C are the canonical Watson-Crick-edge DMS targets, the DMS structural analysis (panels a, b) is restricted to these two positions. Sample sizes are annotated above each bar. The underlying analysis is described in Supplementary Note 1.

#### Supplementary Note

**When reactivity values are projected onto all eight chains of the solved cryo-EM structure of ROOL env-120 (PDB: 9MDS)** (ref 10)**, the three-dimensional distribution of reactivity provides an intuitive visualization of RNA structural features (Fig. 4b). Reactivity is displayed on separate probe-specific color scales (teal for 2A3; coral for DMS), with high-reactivity nucleotides clustering at known flexible regions including the UAA triloop and the interdomain linker, while low-reactivity nucleotides map to the interior of helical stems and tertiary interaction interfaces. Primer-binding regions lie outside the reported region and are shown in lavender (not analyzed).**

Base-pairing assignments for each nucleotide were extracted from the cryo-EM structure (PDB 9MDS) using DSSR (Dissecting the Spatial Structure of RNA), classifying each position as paired (Watson-Crick or non-canonical base pair) or unpaired (loop, bulge, linker, or terminal). Violin plots stratified by structural context (Supplementary Fig. 4a) show that unpaired nucleotides have significantly higher median reactivity than base-paired nucleotides for both probes (Mann-Whitney U test, P values indicated). ROC analysis (Supplementary Fig. 4b) demonstrates that the reactivity profiles can discriminate paired from unpaired nucleotides with high accuracy (AUC: 2A3 all nucleotides = 0.723; DMS A/C only = 0.815). Nucleotide-resolved analysis (Supplementary Fig. 4c) reveals that DMS reactivity protections are clearly visible at positions involved in tertiary contacts, providing structural information beyond secondary structure alone. Reactivity projection onto the 3D structure was performed in ChimeraX by assigning normalized reactivity values as a per-residue attribute (via a define-attribute file) and coloring the structure by that attribute, on a probe-specific gradient (teal for 2A3, coral for DMS), both scaled 0–1.5 to match the two-dimensional panels. The cif-overlay tool (github.com/hmblair/cif-overlay) can automate this projection.

### Supplementary Tables

**Supplementary Table 1 | Oligonucleotide primer sequences used in this protocol.**

| Name | Sequence (5′→3′) | Length (nt) | Tm (°C) | Notes |
|---|---|---|---|---|
| T7GG_F | TTCTAATACGACTCACTATAGG | 22 | 49 | T7 promoter forward primer; used for PCR amplification of the gene fragment and in vitro transcription |
| ROOL_120_FP | GGAATGTTTATAGACATAGC | 20 | 46 | Forward primer for ROOL env-120 amplicon; anneals within the RNA of interest (see CRITICAL STEP at Step 22) |
| ROOL_120_RP | GGAATGTTCACAAATCATAG | 20 | 46 | Reverse primer for ROOL env-120 amplicon; used for both reverse transcription and PCR |

#### Supplementary Data

The archive `FAST-MaP_ROOL-env120_SupplementaryData.zip` accompanies this protocol. It contains the reactivity data shown in the manuscript, the `cmuts` analysis inputs and outputs, the secondary and tertiary structures used for the figures, and the raw pilot sequencing reads.

**`reactivity/` - the data shown in the manuscript**

- `rool120-custom-profiles.csv`, `rool120-pilot-profiles.csv` - per-nucleotide normalized reactivity, standard error, and reads for all four conditions (bicine/HEPES × 2A3/DMS), at custom and pilot depth.
- `per_condition_custom/`, `per_condition_pilot/` - the same data, one CSV per condition (position, nucleotide, reactivity, std_error, reads).
- `reads_SNR_summary.csv` - reads and mean SNR per condition (the Table 3 data).

**`cmuts_inputs/`**

- `rool120-ref.fasta` - reference sequence submitted to `cmuts`.
- `pipeline.sh` - the `cmuts` CLI pipeline used to generate the profiles.

**`cmuts_output/` - re-running `cmuts` on the inputs reproduces these**

- `plasmidsaurus-full_cmuts/` - per-condition mismatch/deletion rate tables (csv/), mutation-count rates (`rates/*.h5`) for all eight custom conditions (including the DMSO and ethanol no-probe controls), and normalized reactivity (`normalized/*.h5`) for the four probe conditions.

**`structures/` - used for the figures**

- `rool120_cryoEM_secondary_structure.dbn` - cryo-EM secondary structure (dot-bracket), used for the 2D overlays and ROC analysis.
- `rool120_forna_layout.json` - FORNA layout for the 2D structure panels.
- `rool120_2A3_reactivity_forna.txt`, `rool120_DMS_reactivity_forna.txt` - per-nucleotide reactivity used to color the 2D overlays.
- `9MDS_cryoEM_tertiary.cif` - cryo-EM tertiary structure (PDB 9MDS; Kretsch et al.), used for the 3D reactivity projections.
- `rool120_sequence.txt` - RNA sequence.

**`raw_sequencing/`**

- `plasmidsaurus-pilot.zip` - raw pilot sequencing (FASTQ) and its `cmuts` run folder.
- The custom high-depth raw FASTQ (~1.7 GB) is not bundled here; it is available from the authors on request.
- `README.md` - full description and provenance (`cmuts` settings, Plasmidsaurus order, numbering).
