## Supplementary material for "FAST-MaP: Chemical Mapping of RNA Structures Using Primer-less Sequencing": FAST-MaP_ROOL-env120_Supplementary_Data: pipeline.sh

```
#!/bin/zsh
# Runs the cmuts pipeline on the rool120 plasmidsaurus data.
# Author: Hamish M. Blair <>

set -e
set -u
set -o pipefail

cd "${0:A:h}"

readonly FASTA=data/ref.fasta
readonly PRESET=map-ont
readonly THREADS=14

readonly ALIGN_DIR=alignments
readonly RATE_DIR=rates
readonly SUB_DIR=subtracted
readonly NORM_DIR=normalized
readonly CSV_DIR=csv

readonly BUFFERS=(hepes bicine)
readonly SAMPLES=(2A3 DMS DMSO ETH)

# The untreated background for each modifier.
background() {
    case "$1" in
        2A3) print DMSO ;;
        DMS) print ETH ;;
    esac
}

make_directories() {
    mkdir -p "$ALIGN_DIR" "$RATE_DIR" "$SUB_DIR" "$NORM_DIR" "$CSV_DIR"
}

align_one() {
    local name="$1"
    cmuts align \
        --fasta "$FASTA" \
        --preset "$PRESET" \
        --threads "$THREADS" \
        --overwrite \
        --output "$ALIGN_DIR/$name.bam" \
        "data/$name.fastq"
}

count_one() {
    local name="$1"
    cmuts hmm \
        --fasta "$FASTA" \
        --workers "$THREADS" \
        --overwrite \
        --output "$RATE_DIR/$name.h5" \
        "$ALIGN_DIR/$name.bam"
}

subtract_one() {
    local name="$1"
    cmuts sub \
        --overwrite \
        --output "$SUB_DIR/$name.h5" \
        "$RATE_DIR/$name.h5" \
        "$RATE_DIR/$2.h5"
}

align_all() {
    for buffer in $BUFFERS; do
        for sample in $SAMPLES; do
            align_one "$buffer-$sample"
        done
    done
}

count_all() {
    for buffer in $BUFFERS; do
        for sample in $SAMPLES; do
            count_one "$buffer-$sample"
        done
    done
}

subtract_all() {
    for buffer in $BUFFERS; do
        for sample in 2A3 DMS; do
            subtract_one "$buffer-$sample" "$buffer-$(background $sample)"
        done
    done
}

# Each condition is normalized on its own, so its scale depends only on its
# own treated and untreated samples.
normalize_one() {
    local name="$1"
    cmuts norm \
        --overwrite \
        --output "$NORM_DIR/$name.h5" \
        "$SUB_DIR/$name.h5"
}

normalize_all() {
    for buffer in $BUFFERS; do
        for sample in 2A3 DMS; do
            normalize_one "$buffer-$sample"
        done
    done
}

write_one_csv() {
    local name="$1"
    cmuts csv \
        --fasta "$FASTA" \
        "$NORM_DIR/$name.h5" \
        > "$CSV_DIR/$name.csv"
}

write_all_csv() {
    for buffer in $BUFFERS; do
        for sample in 2A3 DMS; do
            write_one_csv "$buffer-$sample"
        done
    done
}

main() {
    make_directories
    align_all
    count_all
    subtract_all
    normalize_all
    write_all_csv
}

main
```
