## Supplementary material for "FAST-MaP: Chemical Mapping of RNA Structures Using Primer-less Sequencing": FAST-MaP_ROOL-env120_Supplementary_Data: README.md

# FAST-MaP — ROOL env-120 Supplementary Data

Accompanies the FAST-MaP protocol manuscript. Contains the reactivity data shown in the
manuscript, the cmuts analysis inputs and outputs, the secondary/tertiary structures used
for the figures, and the raw pilot sequencing data.

## Provenance
- Reactivity computed with cmuts ([github.com/DasLab/cmuts](https://github.com/DasLab/cmuts); MIT license), cmuts 2.0.
  Settings: minimap2 `map-ont` alignment, minimum mapping quality 20, background subtraction
  against the paired no-probe control with negative reactivities clipped to zero (the cmuts
  default), and `ubr` normalization applied to each condition independently (per treated/background pair).
  Supplementary/split-read alignments are excluded (cmuts no-supplementary default).
  DMS reactivity = mismatch (substitution) rate; 2A3 reactivity = mismatch + deletion rate;
  insertions are excluded for both probes.
- Reported "reads" = treated + background reads that passed filtering and contributed to the
  profile (primary alignments only; supplementary/split-read records are not counted).
- "custom" = high-depth Plasmidsaurus order (416,000–520,000 reads/condition); "pilot" =
  low-depth pilot order (5,700–10,000 reads/condition). See `reactivity/reads_SNR_summary.csv`.
- Reference = 615 nt (ROOL env-120; transcript positions 21–635 renumbered 1–615).

## Contents

### `reactivity/` — the data shown in the manuscript
- `rool120-custom-profiles.csv`, `rool120-pilot-profiles.csv` : per-nucleotide normalized
  reactivity, standard error, and reads for all four conditions (bicine/HEPES × 2A3/DMS).
- `per_condition_custom/`, `per_condition_pilot/` : the same, one file per condition
  (`position`, `nucleotide`, `reactivity`, `std_error`, `reads`).
- `reads_SNR_summary.csv` : reads and mean SNR per condition (the Table 3 data).

### `cmuts_inputs/` — inputs for the cmuts web server / CLI
- `rool120-ref.fasta` : reference sequence.
- `pipeline.sh` : the cmuts CLI pipeline used to generate the profiles.

### `cmuts_output/` — authoritative cmuts outputs (re-running cmuts on the inputs reproduces these)
- `plasmidsaurus-full_cmuts/csv/` : per-condition normalized reactivity tables (custom, 4 probe conditions).
- `plasmidsaurus-full_cmuts/normalized/` : normalized reactivity (`.h5`), 4 probe conditions.
- `plasmidsaurus-full_cmuts/rates/` : per-nucleotide mutation-count rates (`.h5`), all 8
  custom conditions (incl. DMSO/ETH no-probe controls).

### `structures/` — used for the figures
- `rool120_sequence.txt` : RNA sequence.
- `rool120_cryoEM_secondary_structure.dbn` : cryo-EM secondary structure (dot-bracket),
  used for the 2D overlays and ROC analysis.
- `rool120_forna_layout.json` : FORNA layout for the 2D structure panels.
- `rool120_2A3_reactivity_forna.txt`, `rool120_DMS_reactivity_forna.txt` : per-nucleotide
  reactivity (custom, bicine) used to color the 2D overlays.
- `9MDS_cryoEM_tertiary.cif` : cryo-EM tertiary structure (PDB 9MDS; Kretsch et al.), used
  for the 3D reactivity projections.

### `raw_sequencing/` — raw reads
- `plasmidsaurus-pilot.zip` : raw pilot sequencing (FASTQ) plus its cmuts run folder.
- NOTE: the custom (high-depth) raw FASTQ is ~1.7 GB compressed and is not bundled here;
  it is available from the authors on request.
